# A resource for generating and manipulating human microglial states in vitro

**DOI:** 10.1101/2022.05.02.490100

**Authors:** Michael-John Dolan, Martine Therrien, Saša Jereb, Tushar Kamath, Trevor Atkeson, Samuel E. Marsh, Aleksandrina Goeva, Neal M. Lojek, Sarah Murphy, Cassandra M. White, Julia Joung, Bingxu Liu, Francesco Limone, Kevin Eggan, Nir Hacohen, Bradley E. Bernstein, Christopher K. Glass, Ville Leinonen, Mathew Blurton-Jones, Feng Zhang, Charles B. Epstein, Evan Z. Macosko, Beth Stevens

## Abstract

Microglia have emerged as key players in the pathogenesis of neurodegenerative conditions such as Alzheimer’s disease (AD). In response to CNS stimuli, these cells adopt distinct transcriptional and functional subtypes known as states. However, an understanding of the function of these states has been elusive, especially in human microglia, due to lack of tools to model and manipulate this cell-type. Here, we provide a platform for modeling human microglia transcriptional states *in vitro*. Using single-cell RNA sequencing, we found that exposure of human stem-cell differentiated microglia (iMGLs) to brain-related challenges generated extensive transcriptional diversity which mapped to gene signatures identified in human brain microglia. We identified two *in vitro* transcriptional clusters that were analogous to human and mouse disease-associated microglia (DAMs), a state enriched in neurodegenerative disease contexts. To facilitate scalable functional analyses, we established a lentiviral approach enabling broad and highly efficient genetic transduction of microglia *in vitro*. Using this new technology, we demonstrated that MITF (Melanocyte Inducing Transcription Factor), an AD-enriched transcription factor in microglia, drives both a disease-associated transcriptional signature and a highly phagocytic state. Finally, we confirmed these results across iMGLs differentiated from multiple iPSC lines demonstrating the broad utility of this platform. Together, these tools provide a comprehensive resource that enables the manipulation and functional interrogation of human microglial states in both homeostatic and disease-relevant contexts.

## Introduction

Microglia, the macrophages of the brain, perform central roles in development, homeostasis and diseases of the central nervous system (CNS)^1, 2^. Recent work has highlighted the key role of this cell type in neurodegenerative disorders such as Alzheimer’s disease (AD). Emerging genetic studies of late onset AD implicate microglia, with a large fraction of AD risk genes specifically expressed in myeloid cells^3, 4^, while microglia activation and dysfunction is a hallmark of AD and other neurodegenerative disorders^5^. The application of single-cell RNA sequencing (scRNAseq) has revealed that microglia exhibit multiple states^6–14^, subpopulations that express distinct gene signatures, including disease-associated microglia (DAMs) which occur in neurodegenerative disorders. However, the function of these states and their impact on disease remains unknown.

Since microglial states are defined by the differential expression of many genes, scalable models are needed to understand the function of transcriptional signatures. However, there are several outstanding challenges: Many differences exist between mouse and human microglia^15, 16^, a problem which is particularly acute in the study of AD risk genes^17^. Microglia also rapidly alter their gene expression upon primary culture, presumably due to a lack of brain-derived cues^15, 18^ while cell lines fail to accurately model this cell-type^19^. Finally, high-throughput gene perturbation studies are not possible as this cell-type is resistant to most conventional forms of DNA delivery^20^.

Here, using human stem cell-derived microglia (iMGLs), we address these issues by creating an *in vitro* platform for studying microglial states in human cells. By exposing iMGLs to brain-related substrates and performing scRNAseq and cross-dataset integration, we show that many transcriptional states identified in the human brain can be recapitulated in a monolayer culture. In particular, we identify DAM-like states *in vitro* and demonstrate their formation is dependent on Triggering Receptor Expressed on Myeloid cell 2 (TREM2) signaling, as has been observed *in vivo*^7, 21^.

To enable scalable manipulations of human microglia, we established a lentiviral transduction protocol that opens up iMGLs to viral manipulation and large-scale screens. We use this new transduction approach to determine that overexpression of Melanocyte Inducing Transcription Factor (MITF), a transcription factor upregulated in microglia of AD patients, drives disease-associated genes and a highly phagocytic state. Finally, we demonstrate that our observations are robust by differentiating iMGLs from multiple human induced pluripotent stem cell (iPSC) lines, enabling investigators to apply our tools to patient-derived stem cells. The datasets and approaches provided here will enable a systematic dissection of human microglial states and are applicable to the study of this cell-type not just disease, but throughout development, homeostasis and repair.

## Results

### CNS substrates induce transcriptional states in iMGLs analogous to those found in the human brain

Microglia cultured *in vitro* lack many of the key transcriptional features observed *in vivo*^15, 18^. We hypothesized that exposure to complex brain substrates or cellular debris may drive the formation of brain-like microglial states in a dish. To test this, we differentiated human embryonic stem cells into iMGLs^22^ and at day 40 we performed single cell RNA sequencing (scRNAseq) on cells either untreated or challenged with synaptosomes, myelin debris, apoptotic neurons or synthetic amyloid-beta (Aβ) fibrils (Fig. 1A). We acquired a total of 57,837 single iMGL transcriptomes across all conditions (Fig. 1B, Fig. S1A-D, Supplementary Table 1) and cells showed enrichment of microglial identity genes with very little expression of pluripotency markers (Fig. S1E-F), confirming successful differentiation.

**Figure 1:**
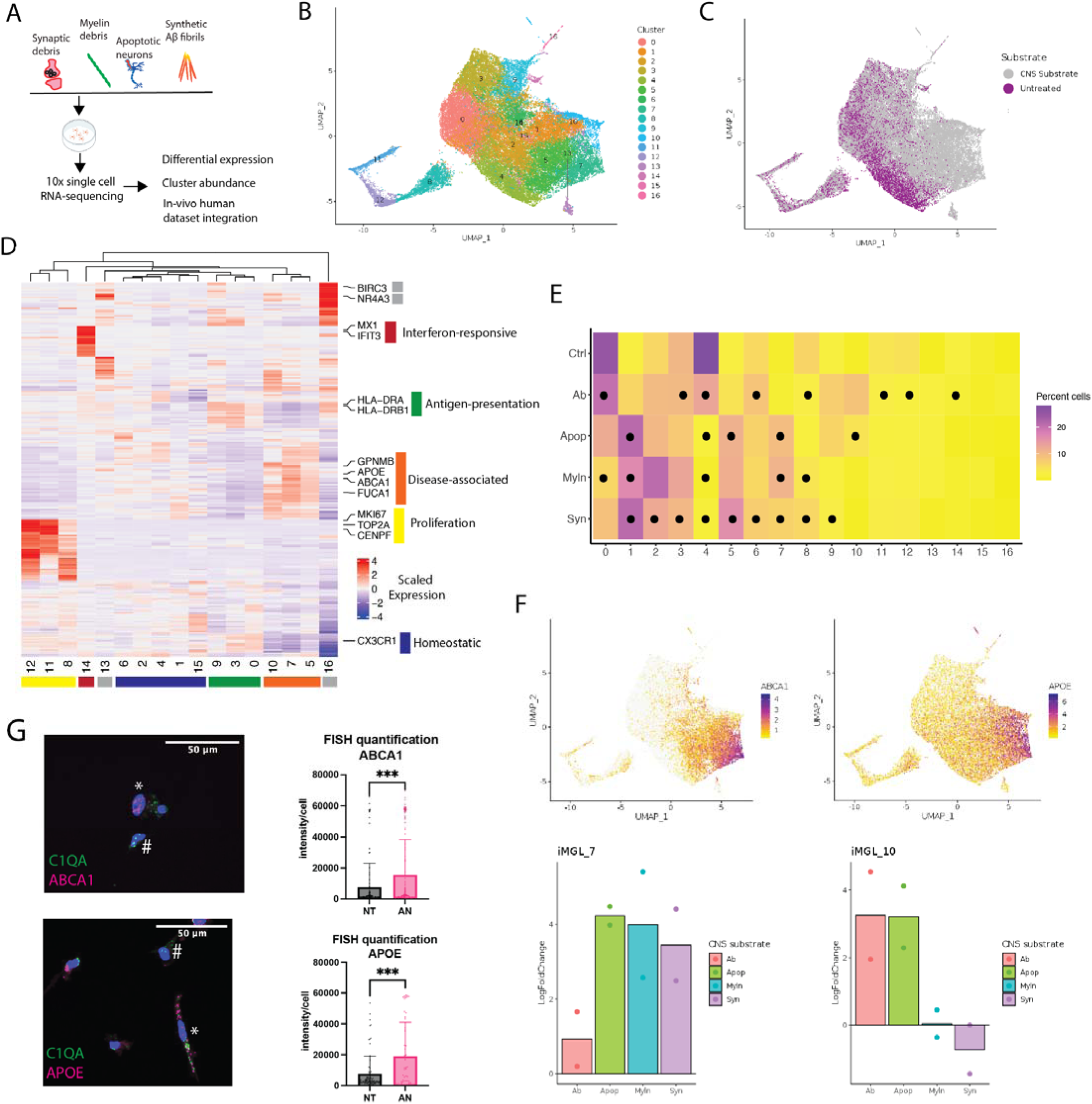
Treatment of iMGLs with CNS challenges induces diverse transcriptional states that map to those found *in vivo*. A) iMGLs were either untreated or treated for 24h with synaptosomes, myelin debris, synthetic Aβ fibrils or apoptotic neurons (collectively CNS substrates) followed by scRNAseq. B) Uniform Manifold Approximation and Projection (UMAP) projection of iMGLs; total of 57,837 cells, cells colored by cluster. C) UMAP projection, cells colored by untreated or CNS substrate treated condition. D) Heatmap of top 20 most differentially enriched genes for each cluster, see Fig. S4A for all differentially expressed genes. Clusters are sorted by similarity and larger groups of clusters labeled. E) Mean relative abundance of each cluster across each condition. Circle represents significant difference (ie. adjusted p-value<0.05) determined by a Dirichlet regression test for differential abundance. Syn=synaptosomes, Myln=myelin debris, Ab=synthetic Aβ fibrils, Apop= apoptotic neurons. F) Marker gene expression (top) and log fold change of cluster relative to untreated (bottom) for iMGL_7 (left) and iMGL_10 (right) (Syn=synaptosomes, Myln=myelin debris, Ab=synthetic Aβ fibrils, Apop= apoptotic neurons). G) Validation of gene expression with fluorescent in-situ hybridization (FISH), showing enrichment of markers for iMGL_7 and iMGL_10 in the presence of apoptotic neurons using smFISH (NT= non tre ted, AN= apoptotic neurons). Top, representative image showing C1QA (green) and ABCA1 (magenta) mRNA. Bottom, C1QA (green) and APOE (magenta) mRNA. Right, quantitative analysis of marker gene enrichment (iMGL_7: ABCA1 *** p-value = 0.0006) and (iMGL_10: APOE ***p-value = 0.0001). Error bars represent standard deviation. Symbol * indicates a positive cell and # indicate a negative cell.

Using unbiased clustering, we identified 17 different clusters of iMGLs which were persistent across replicates (Fig. 1B and Fig. S1D), demonstrating a surprisingly degree of transcriptional diversity as found in a large dataset of human postmortem microglia^12, 23–25^. Exposure to CNS challenges induced diversification of iMGL clusters (Fig. 1C and Fig. S2). Importantly, these clusters were not a general signature of phagocytosis, as each substrate drove distinct gene programs (Fig. S3A) and although all cells phagocytosed substrates (Fig. S3B), not all cells adopted CNS substrate-induced states (see below). This demonstrates that iMGLs, even in a monoculture, are not a homogeneous population but can form diverse transcriptional states in response to different brain substrates or challenges, as observed *in vivo*^1, 2^.

To understand the biological significance of these clusters, we performed a differential expression analysis (Fig. 1D and Fig S4A, see Supplementary Table 2 for full list). Based on gene signatures, the majority of the iMGL clusters (15/17) could be grouped into established microglial states, analogous to those found *in vivo*^26^. We identified cell states similar to those implicated in the neurodegenerative DAM response (clusters iMGL_7 and iMGL_10, expression of *APOE*, *GPNMB*, *LPL* and *ABCA1*), antigen-presentation (clusters iMGL_9, iMGL_3, expression of *HLA-DRA* and *HLA-DRB1*), interferon-responsivity (cluster iMGL_14, expression of *IFIT3* and *MX1*), proliferation (clusters iMGL_8, iMGL_11 and iMGL_12 and expression of *MKI67* and *TOP2A*) and a homeostatic state (expression of *CXCR1*) (Fig. 1D and Fig. S4C-F). These transcriptional signatures have been identified in both mouse and human scRNAseq datasets^6–8, 10–13, 23, 24, 26^, suggesting iMGLs can adopt *in vivo*-like transcriptional states in a dish (see below for dataset integration). The remaining two unknown clusters (iMGL_13 and iMGL_16) consisted of few cells (1.9% of iMGLs) and exhibited high expression of an artifactual activation gene signature (Fig. S4B), likely due to handling^27^. Apart from these two clusters, our dataset was devoid of this activation signature.

We next asked whether exposing iMGLs to specific brain-relevant substrates resulted in context-dependent cell states. We observed many statistically significant changes in iMGL cluster proportions after CNS substrate exposure (Fig. 1E and Fig. S5A, Supplementary Table 3). A few clusters were broadly enriched or depleted in response to multiple CNS challenges (Fig. 1E and Fig. S5B-C). However, we also observed clusters enriched for specific substrates, in particular clusters iMGL_7 and iMGL_10. Cluster iMGL_7 was broadly induced by all substrates except Aβ fibrils, while iMGL_10 was only observed in the presence of apoptotic neurons and Aβ fibrils (Fig. 1E-F). We validated these findings with fluorescent *in situ* hybridization for cluster-enriched markers, confirming the enrichment of iMGL_7 and iMGL_10 in response to apoptotic neurons (Fig. 1G) and the existence of proliferating and interferon-response iMGLs (Fig. S4E’ and S4F’). These results validate our transcriptomic observations and confirm that iMGLs form diverse cell states with distinct transcriptional profiles in response to CNS stimuli, providing a resource for *in vitro* experiments targeting these specific cell states.

We next sought to systematically compare our iMGL transcriptional states with those found *in vivo*, using a single nucleus RNA sequencing (snRNA-seq) dataset from human cortical biopsies of 58 subjects (73,267 single microglial nuclei, Kamath *et al*. in preparation, Fig. S6). To compare iMGL and human brain datasets we used LIGER^28^, which identifies dataset-specific and shared gene sets (metagenes) to integrate and cluster the two datasets jointly (Fig. 2A). Integrative analysis of the human brain and iMGL profiles using LIGER (with default parameters) produced a set of joint clusters with significant alignment between the two datasets (Fig. 2B and S7A). To quantify the alignment we calculated, for each cell, the degree to which its neighbors were from one or all datasets (0 or 1 respectively)^28^. This analysis revealed good mixing between *in vivo* microglia and iMGLs (alignment score=0.76). Microglia from human patients and iMGLs contributed to the majority of joint clusters (Fig. 2B-D, S7B) and we identified conserved states, such as interferon responsive and disease-associated, by an alignment-independent enrichment analysis using marker gene sets (Fig. S7C, see S7D-G for example genes). These results confirm that iMGLs can exhibit transcriptional profiles similar to those found in human brain microglia^26^.

**Figure 2:**
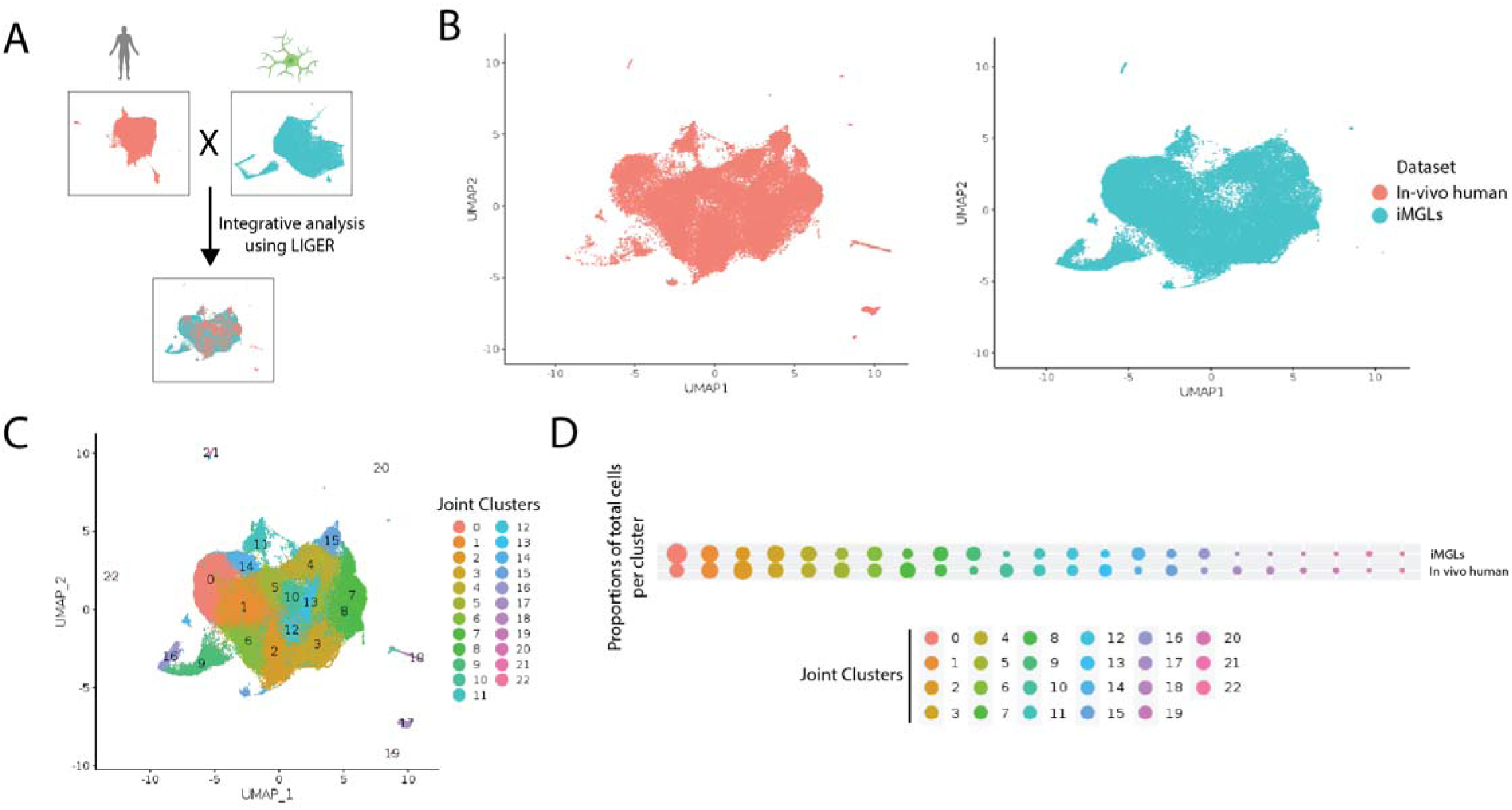
Dataset integration of iMGL and in vivo human brain microglia. A) Schematic showing integration of scRNAseq data using LIGER B) UMAP projection of integrated *in vivo* human microglia (left) and iMGL profiles (right) show good concordance and few groups of dataset specific cells C) Joint clustering resulting from the LIGER integrative analysis. D) Proportion of cells per joint cluster for each source of data (*in vivo* human microglia or iMGLs), dots are sized by proportion of total cells per dataset.

Exposure to CNS substrates drove iMGL transcriptional diversity and disease-associated gene expression across the joint clusters (Fig. S7H), suggesting that addition of CNS substrate is key to producing many in vivo-relevant iMGL clusters. To further test the impact of substrate exposure on human brain-like transcriptional states, we performed the same dataset integration using only untreated iMGLs which led to a poorer alignment with many large clusters populated by human microglia only (Fig. S7I, alignment score=6.25). This demonstrates that CNS substrate challenge enables the generation of several human brain-like transcriptional states in a dish, including neurodegenerative-associated signatures, opening up these states to functional experiments and mechanistic studies.

### Induction of the neurodegenerative DAM microglial state *in vitro*

To further validate the *in vitro* microglial states generated after CNS substrate exposure, we focused on the DAM-like iMGL clusters as this transcriptional signature is highly enriched in neurodegeneration ^7, 21, 23, 24, 29–32^. DAMs are enriched in both human AD patients^23, 24^ and mouse models of neurodegeneration^7, 29^ and this signature has been shown to be dependent on TREM2 receptor signaling^7, 21, 25, 29^. To confirm the existence of this state in our *in vitro* platform, we compared several established DAM datasets to iMGLs and tested the dependency of this *in vitro* state on TREM2 (Fig. 3A).

**Figure 3:**
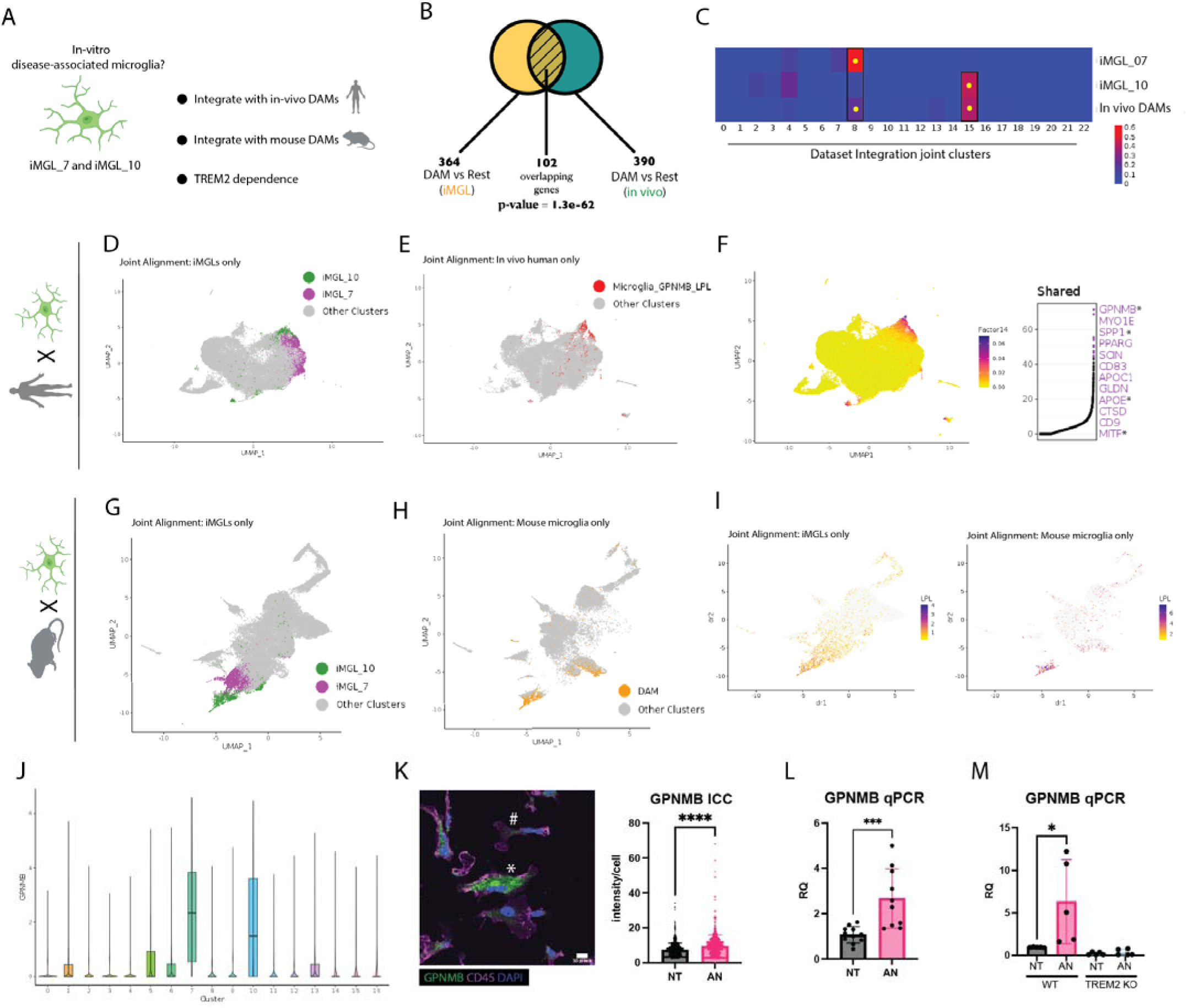
Disease-associated microglial states generated *in-vitro* are similar to those found in human AD and mouse amyloid models. A) Schematic of analyses and experiments used to validate *in vitro* DAM states. B) A differential expression test without dataset integration. Calculated by comparing DAMs within each dataset (iMGL and *in vivo* human microglia) to other clusters from the same dataset, identified 102 overlapping genes (and this overlap was statistically significant: hypergeometric test). C) Percentage distribution of cells from iMGLs (iMGL_7 and iMGL_10) and *in vivo* human (Microglia_GPNMB_LPL, DAMs) over all joint clusters from the LIGER integration. A yellow circle indicates statistically significant enrichment (Binomial test). D) LIGER dataset integration of *in vivo* human brain microglia and iMGLs, showing only iMGLs with cluster iMGL_7 (green) and iMGL_10 (magenta) highlighted. E) *In vivo* human and iMGL LIGER integration showing only human microglia (gray) with human DAMs (Microglia_GPNMB_LPL) labeled in red. F) Left, UMAP projection of LIGER integration showing the shared metagene common to both iMGL and *in vivo* DAMs. Right, top constituent genes of this shared factor. Symbol * labels genes highlighted elsewhere in this study. G) LIGER dataset integration of mouse microglia from an AD model (5xFAD) showing only iMGLs and highlighting cluster iMGL_7 (green) and iMGL_10 (magenta). H) LIGER dataset integration of iMGLs and mouse microglia from an AD model (5xFAD) showing only mouse microglia with DAMs labeled (orange). I) UMAP projection of LIGER integration showing expression of *LPL*, a DAM marker gene. Left, in iMGLs and right, in mouse microglia. J) Violin plot of GPNMB mRNA expression in the iMGL scRNAseq dataset plotted by the iMGL cluster. Highest expression observed in iMGL_7 and iMGL_10. K) Validation of GPNMB protein expression in the presence of apoptotic neurons by immunocytochemistry. Left, representative image showing GPNMB (green) and CD45 (magenta, microglia/macrophage marker). Symbol * indicates a positive cell and # indicate a negative cell. Right, quantitative analysis of GPNMB protein levels (****p-value < 0.0001).Error bars represent standard deviation. L) Validation of GPNMB expression by RT-qPCR when iMGLs are treated with apoptotic neurons (p-value= 0.0009) Error bars represent standard deviation. M) Induction of GPNMB in the presence of apoptotic neurons also observed in an EGFP-expressing cell line (p-value = 0.0427) but not in an isogenic TREM2 knockout. For all RT-qPCR experiments biological replicates are shown. NT=not treated, AN=apoptotic neurons. Error bars represent standard deviation

To further compare DAM-like iMGL and *in vivo* DAMs observed in the human brain (Kamath et al in preparation), we first performed two alignment-independent comparisons of the two DAM signatures. We first calculated two “within dataset” differentially expressed DAM gene lists by comparing DAM-like iMGL (clusters iMGL7 and iMGL10 signature combined) and *in vivo* DAMs (Microglia_GPNMB_LPL) to all other clusters from their respective datasets (Fig. 3B). Comparison of these gene lists confirmed a significant shared transcriptional signature between DAM-like iMGLs and *in vivo* DAMs (Fig. 2B, p-value=1.3 x 10^-62^, Supplementary Table 4). We further validated this by performing a gene set enrichment analysis of human DAM marker genes across iMGL clusters (Fig. S8), and found genes specific to DAMs that were demonstrated to be enriched in AD patient brains and colocalize with amyloid plaques^24^.

In our iMGL and *in vivo* human dataset integration (Fig. 2), we observed that iMGL_7 and iMGL_10 cells were significantly enriched in the same joint clusters as *in vivo* DAMs (Microglia_GPNMB_LPL) (Fig. 3C-E). We also found these cells share a transcriptional signature enriched for genes highly relevant for neurodegeneration, such as *GPNMB*, *SPP1*, *APOE* and *CD9* (Fig. 3F, see Supplementary Table 5 for full shared gene list)^7, 23, 24^. These results demonstrate that CNS-substrate exposure can induce the DAM transcriptional state in a dish.

To further compare the iMGL DAM signature to the *in vivo* microglial state, we integrated iMGL scRNAseq data with mouse and xenotransplanted human microglia exposed to amyloid plaques in an AD mouse model. Integration of the iMGL dataset with mouse 5xFAD microglia^7^ (Fig. 3G-I and Fig. S9A-D) and xenotransplanted human iMGLs^33^ (Fig. S9E-K), revealed extensive alignment of iMGL DAMs, especially iMGL_10, and highlighted shared common gene signatures between mouse and human Aβ-induced DAMs (Fig. 3I and Fig. S9D). These analyses further confirm the existence of a DAM-like state *in vitro* and highlight the key role Aβ plays in DAM formation.

Previous work revealed that mouse and human DAMs fall on a gene expression continuum^7, 24^ and we wondered if this could explain the existence of multiple DAM-like clusters in iMGLs. We ordered apoptotic neuron-exposed iMGLs (which included all clusters, 12,217 cells, n=2) on a pseudotime trajectory based on the similarity of their transcriptional profiles^34^. This analysis revealed a transition from clusters iMGL_5 to iMGL_7 to iMGL_10 (Fig. S10), suggesting that CNS challenges differentially drive iMGLs along a disease-associated trajectory and explaining the existence of multiple DAM-like iMGL clusters.

A critical feature of both mouse and human DAMs is their dependence on TREM2 receptor signaling to form disease-associated signatures^7, 21, 29^. TREM2 was highly expressed in iMGL_7 and iMGL_10 and elevated when cells were treated with apoptotic neurons (Fig. S11A). We identified *Glycoprotein NMB* (*GPNMB)* as a marker gene that specifically labels both DAM clusters in iMGLs (Fig. 3J) and is also enriched in this microglial state *in vivo* (Fig S6E and Fig. 3F). GPNMB was also previously identified as a marker for DAMs^7, 21, 23, 33^. Using immunocytochemistry and real-time quantitative PCR (RT-qPCR), we confirmed that apoptotic neuron exposure drives an increase in GPNMB protein and mRNA (Fig. 3K-L) in iMGLs. Using a previously characterized iPSC line in which TREM2 was deleted and its isogenic control^21^, we found that this increase in *GPNMB* was abolished when TREM2 was absent (Fig. 3M), functionally confirming the identity of the *in vitro* DAM-like state in iMGLs.

The generation of human DAM microglia *in vitro* provides a platform for probing the regulation of this transcriptional state and its functional relevance. The expression of *GPNMB* was dependent on phagocytosis, as blockade of engulfment by cytochalasin D inhibited expression (Fig. S11B). However, DAM formation was specific to CNS substrates, as *GPNMB* was not induced by presence of *E. Coli* (Fig. S11C). Together, these data suggest that disease-associated states *in vitro* share characteristics with human brain and mouse DAMs, and are formed specifically in response to CNS challenges.

### A lentiviral system for highly efficient genetic modification of iMGLs

The ability to model human microglial states in a dish is the first step in enabling functional dissection. However, the necessary scalable approaches to genetically manipulate microglia do not exist, as microglia are resistant to various forms of DNA delivery, such as transfection, electroporation and viral transduction^20^. To augment the iMGL platform for functional studies, we sought to develop a robust lentiviral technology capable of transducing microglia. Lentivirus have a large genetic capacity and regularly used as a vector for genetic screens^35^.

Previously, successful transduction of monocytes and monocyte-derived dendritic cells with HIV and lentivirus was achieved using a protein from simian immunodeficiency virus, Vpx^36–38^. Vpx degrades SAMHD1, a restriction factor present in human cells that degrades dNTPs, which in turn prevents efficient reverse transcription of lentiviral RNA^39, 40^. We found that co-delivery of Vpx packaged in VSV-G virus-like particles (VLPs) dramatically improves lentiviral transduction of iMGLs, only ∼4% of iMGLs were transduced with lentivirus alone. However, transduction efficiency increased to 89% when Vpx VLPs were added (Figure 4A). Furthermore, Vpx VLPs improved transduction of iMGLs derived from five different iPSC lines (Fig. S12A), confirming Vpx has a robust effect regardless of iMGL source.

**Figure 4:**
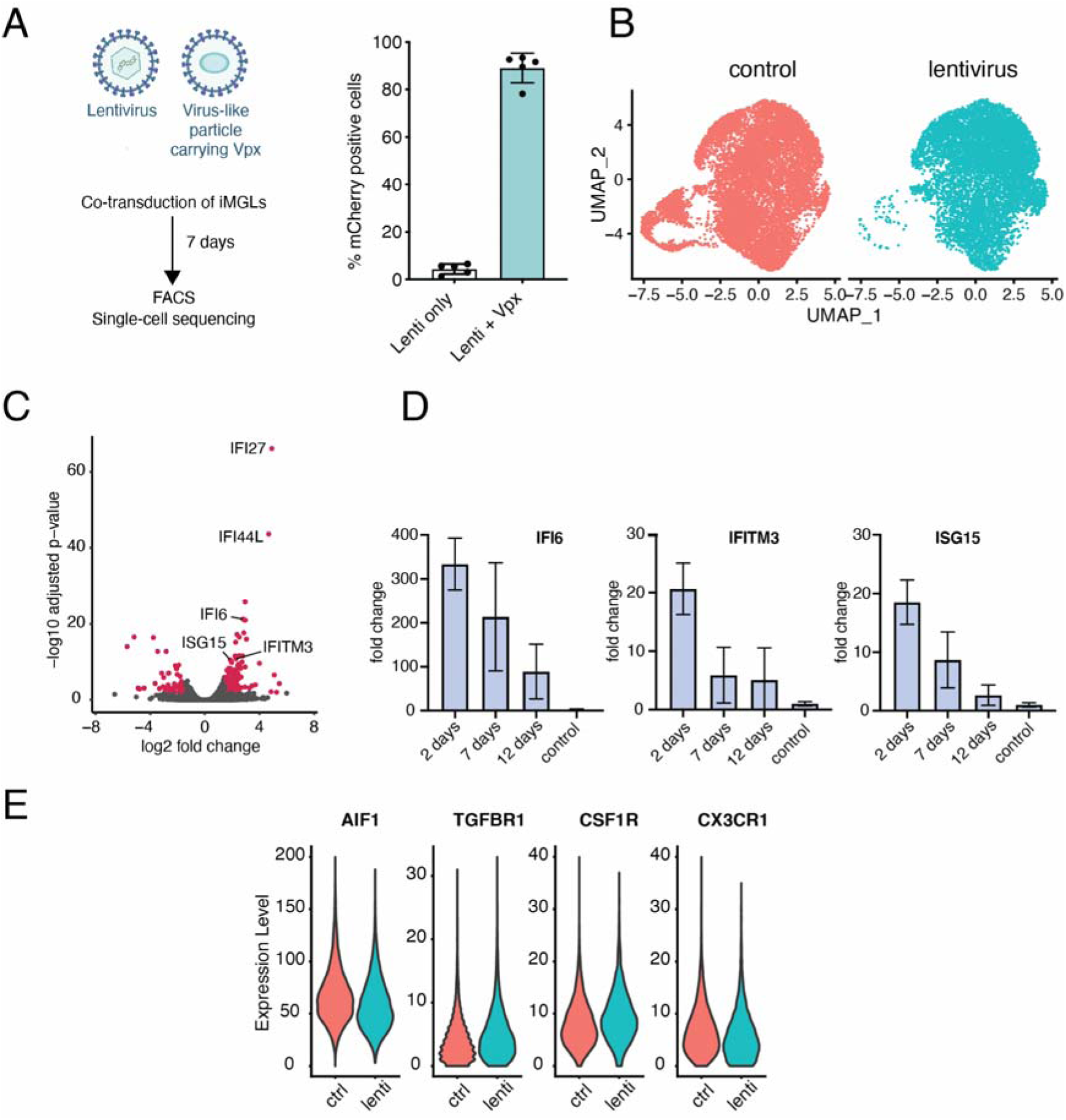
Lentivirus-mediated genetic modification of iMGLs. A) Transduction efficiency of iMGLs with lentivirus only and l ntivirus with VPX virus-like particles, determined by flow cytometry (p-value = 0.0079, two-tailed Mann-Whitney test). B) UMAP projection of single cell sequencing data from two control and lentivirus transduced iMGL differentiations. C) Volcano plot of differentially-expressed genes between control and lentivirus transduced samples. Genes highlighted in red have adjusted p-value < 0.001 and log2 fold change < 2. D) RT-qPCR time-course analysis of expression of three interferon-induced genes in iMGLs after lentivirus transduction. E) Expression of microglial markers in the control and lentivirus samples. P-values for comparisons between control (ctrl) and lentivirus (lenti) are not significant for all genes shown (AIF1 p-value=0.463, TGFBR1 p-value=0.999, CSF1R p-value=0.999 and CX3CR1 p-value=0.2193, Wald test with multiple hypothesis correction was performed with DESeq2).

Lentiviral exposure induces transcriptional response in many cell-types, especially cells of the immune system^41^, which could confound experimental interpretation. To determine the impact of lentiviral exposure on iMGLs, we co-transduced cells with lentivirus and Vpx VLPs. Seven days after transduction, we performed scRNAseq on transduced and control cells. We found control and lentivirus treated cells clustered together (Fig. 4B), though lentivirus-transduced samples had on average fewer proliferating cells compared to controls (24.8% in controls vs. 11.9% in lentivirus treated samples, Fig. S12B). In addition, expression of 156 interferon-stimulated genes was significantly increased in lentivirus-transduced samples compared to controls (significantly increased genes were defined as those with absolute log2 fold change >= 1.5 and adjusted p-value < 0.01; Fig. 4C, S12C and Supplementary Table 6). However, a RT-qPCR of a representative subset of known interferon-stimulated genes showed downregulation over time (Fig. 4D). In addition, lentivirus treatment did not affect expression of key microglia markers (*AIF1, TGFBR1, CSF1R, CX3CR1;* Fig. 4E). Therefore, when combined with Vpx VLPs, lentivirus can be used as an efficient tool to manipulate gene expression in iMGLs while minimally affecting the iMGL transcriptional profile.

### Identification of MITF as a driver of disease-associated microglia signature and phagocytic function

The ability to virally manipulate iMGLs provided an opportunity to probe the function of genes enriched in the DAM transcriptional state. We chose to focus on transcription factors as these are likely key regulators of changes in cellular function^42^. To identify key regulators of DAM, we combined epigenetic and gene regulatory network analyses (Fig. 5A). First, to identify changes in chromatin accessibility upon DAM formation, we performed an assay for transposase-accessibility chromatin using sequencing (ATAC-seq) on either untreated or apoptotic neuron-exposed iMGLs. Treatment of iMGLs with apoptotic neurons increased the number of differentially accessible chromatin regions (DAR) compared to control (Fig. S13A-C). HOMER^43^ was then used to identify enriched motifs in the regions with increased accessibility following treatment with apoptotic neurons, nominating potential transcription factors regulating gene expression in response to apoptotic neurons (Supplementary Table 7). This demonstrates exposure to brain challenges induce extensive changes in chromatin accessibility in iMGLs.

**Figure 5:**
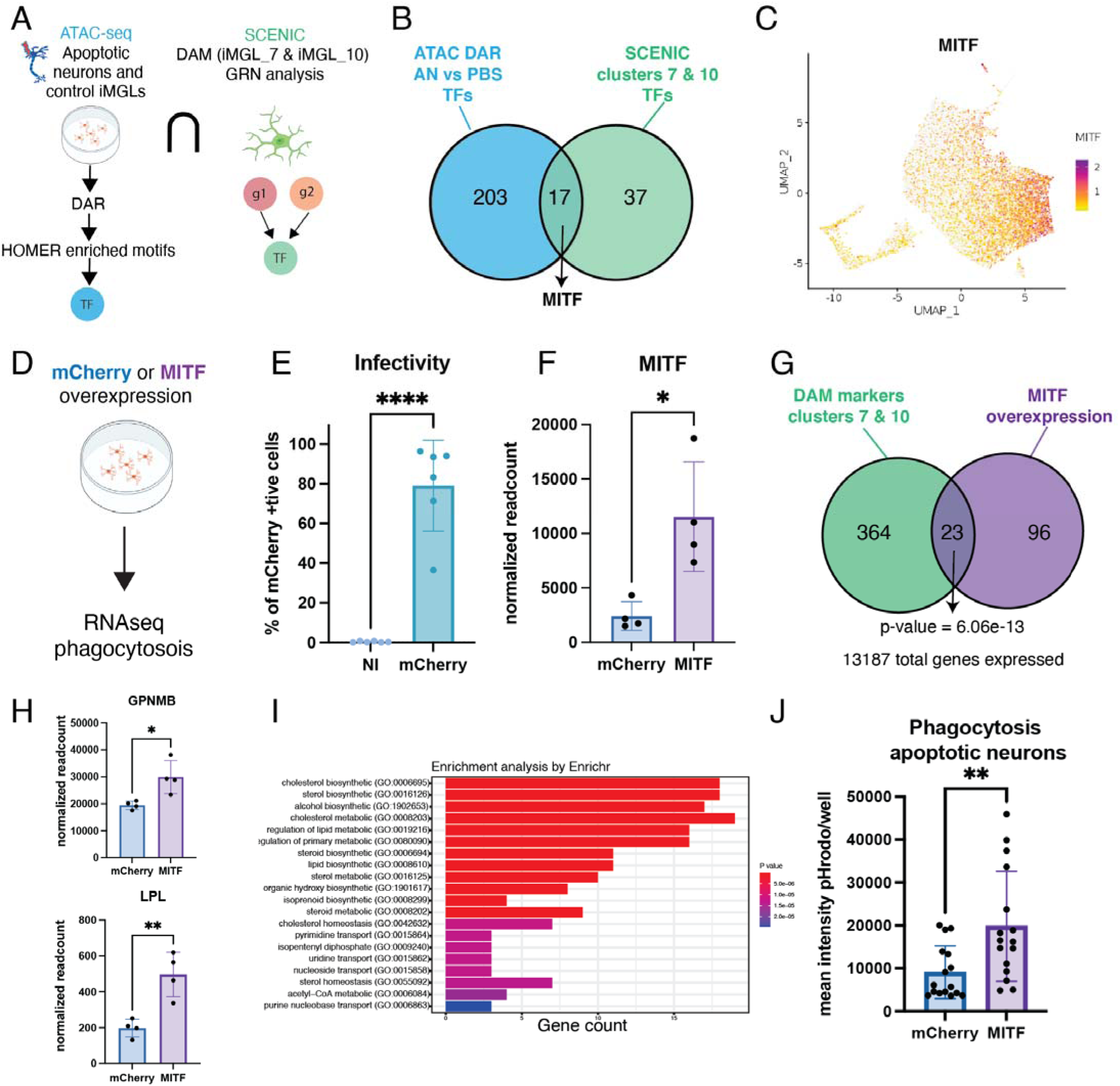
Identification of MITF as a key DAM regulator and driver of phagocytosis. A) Schematic of experiments performed to analyze the effect of apoptotic neurons on iMGL chromatin accessibility to identify differentially accessible chromatin regions (DAR), gene regulatory networks (GRN). B) Intersection of transcription factors (TFs) nominated by ATAC-seq and SCENIC analysis. C) UMAP projection showing MITF expression in the iMGL dataset shows enrichment in DAM clusters. D) Schematic of MITF overexpression experiment, a lentivirus expressing mCherry or MITF was introduced into iMGLs followed by transcriptional (bulk RNA-seq) or functional (phagocytosis) assays. E) Percentage of mCherry cells determined by flow cytometry highlights the number of cells transduced successfully (p-value< 0.0001) (NI= non-infected).Error bars represent standard deviation. F) Quantification of MITF expression in iMGLs transduced with mCherry- and MITF-expressing lentivirus (p-value = 0.0129).Error bars represent standard deviation. G) Venn diagram of overlap between genes differentially expressed between iMGL samples transduced with MITF- expressing versus mCherry-expressing lentivirus and genes differentially expressed in iMGL_07 and iMGL_10 (Figure 1D). P-value is calculated using a hypergeometric test. H) Top, quantification of *GPNMB* expression in RNA-seq samples (p-value=0.0159).Error bars represent standard deviation. Bottom, quantification of *LPL* expression in RNA-seq samples (p-value=0.0042). I) Gene Ontology analysis of genes enriched in MITF-overexpressing iMGLs reveals enrichment of cholesterol and lipid metabolism biological processes. Both length of bar and shading represent relative statistical significance. J) Phagocytosis of pHrodo-conjugated apoptotic neurons in iMGLs transduced with either MITF and mCherry, mean intensity per well measured by flow cytometry (p-value= 0.0051). Error bars represent standard deviation

To identify potential regulators of specific microglial states, we next performed a gene regulatory network analysis on iMGL single-cell profiles using SCENIC^44^, identifying differential expression of transcription factors and their putative direct targets (together termed regulons) (Fig. S13D-E, see Supplementary Table 8 for analysis on all iMGL clusters). We examined the most enriched regulatory networks in clusters iMGL_5, iMGL_7 and iMGL_10 (Fig. S13D-E), and while many regulons are shared across these 3 clusters (eg. CEBPD, IRF8), we also identified several that were differentially expressed in each cluster (eg. ATF3 regulon in iMGL_10, NFE2L2 regulon in iMGL_7 and iMGL_10). Although we focus on DAM iMGLs, this analysis revealed potential regulatory networks underlying all iMGLs clusters (see Supplementary Table 8).

Intersection of transcription factors nominated by both chromatin accessibility and SCENIC analyses (Fig. 5A) revealed 17 putative regulators of the DAM state (Fig. 5B, Supplementary Table 7), including MITF, MAFB and JUN among others. We focused our efforts on MITF, which has recently been identified as upregulated in microglia of patients with AD ^23, 24^ and nominated as a regulator of the neurodegenerative signature in a recent meta-analysis^45^. Correspondingly, MITF mRNA is highly upregulated in DAM-like clusters iMGL_7 and iMGL_10 (Fig. 5C) and its expression scales with the disease-associated trajectory identified in iMGLs (Fig. S10). However, the role of this transcription factor in microglial function is unknown.

To understand the impact of MITF on the iMGLs gene expression and function, we leveraged our iMGL lentiviral approach to overexpress this transcription factor in iMGLs (Fig. 5D). We transduced iMGLs with a lentiviral vector expressing either MITF or mCherry as a control and seven days post infection we performed bulk RNA-seq to determine gene expression changes. On average, 79% of iMGLs were transduced with a mCherry control vector (Fig. 5E). Transduction of iMGLs with MITF cDNA induced differential expression of 670 genes compared to the mCherry control (FDR<0.05, 300 upregulated genes, 371 downregulated genes, Fig. S14A-B, see Supplementary Table 9), including *MITF* itself, confirming overexpression (Fig. 5F). Strikingly, we observed significant overlap between genes upregulated in iMGL DAMs and those driven by MITF (iMGL_7 and iMGL_10, 23 genes p-value = 6.06 x 10^-13^, Fig. 5G, Supplementary Table 9), including DAM marker genes *GPNMB* and *LPL* (Fig. 5H). Finally, a gene ontology analysis of MITF-overexpressing iMGLs showed an enrichment of genes involved in cytokine signaling and cholesterol metabolism (Fig. 5I), pathways found to be upregulated in the DAM transcriptional signature^7, 23, 24, 29^.

What are the functional consequences of MITF upregulation? An increase in phagocytosis, a key microglial function, has been described in diverse contexts including neurodegenerative disease models^5^. To test if MITF plays a role in regulating phagocytosis, we exposed mCherry- or MITF-transduced iMGLs to pHrodo-labeled apoptotic neurons. Strikingly, iMGLs overexpressing MITF exhibited increased phagocytosis compared to controls (Fig. 5J), suggesting that this DAM regulator plays a role in inducing a highly phagocytic state in microglia. Collectively, these results suggest that MITF is an important regulator of the neurodegenerative disease-associated signature including stimulating increased phagocytic function^23, 24, 45^.

### Microglia derived from multiple iPSC lines exhibit transcriptional diversity and DAM induction

A key advantage of stem-cell derived models is the ability to perform parallel experiments on iPSC lines derived from multiple patients^46^. To confirm that our findings were broadly applicable to iMGLs differentiated from multiple donors, we differentiated iMGLs from three independent iPSC lines derived from healthy individuals (all lines were APOE 3/3 status with no TREM2 mutation, see methods for details), exposed these cells to apoptotic neurons (or untreated control) and performed scRNAseq (Fig. 6A). To dissect the impact of iPSC lines on iMGL states, we integrated the two iMGL transcriptomic datasets (H1-derived and iPSC-derived iMGLs) using LIGER (Fig. 6A-C, Supplementary Table 10). This revealed a high level of similarity between the two datasets (alignment score=0.973, Fig. 6B and Fig. S15A-C) and further analysis revealed similar transcriptional states as those identified in iMGLs previously (Fig. S16). Finally, we tested if exposure to apoptotic neurons drives the DAM signature observed in H1-derived iMGLs above. In this analysis both iMGL_7 and iMGL_10 mapped to joint cluster 7 of the integrated iMGL dataset (Fig. 6C-E) which corresponded to a specific metagene consisting of DAM signature genes (Fig. 6D) including expression of *GPNMB* and *LPL* (Fig. S17). Critically, iPSC-derived iMGLs from joint cluster 7 were significantly enriched in the presence of apoptotic neurons (Fig. 6F, p-value = 0.00471), demonstrating that microglia differentiated from iPSCs also exhibit the DAM state in the presence of apoptotic neurons and highlights the generalized nature of our iMGL platform.

**Figure 6:**
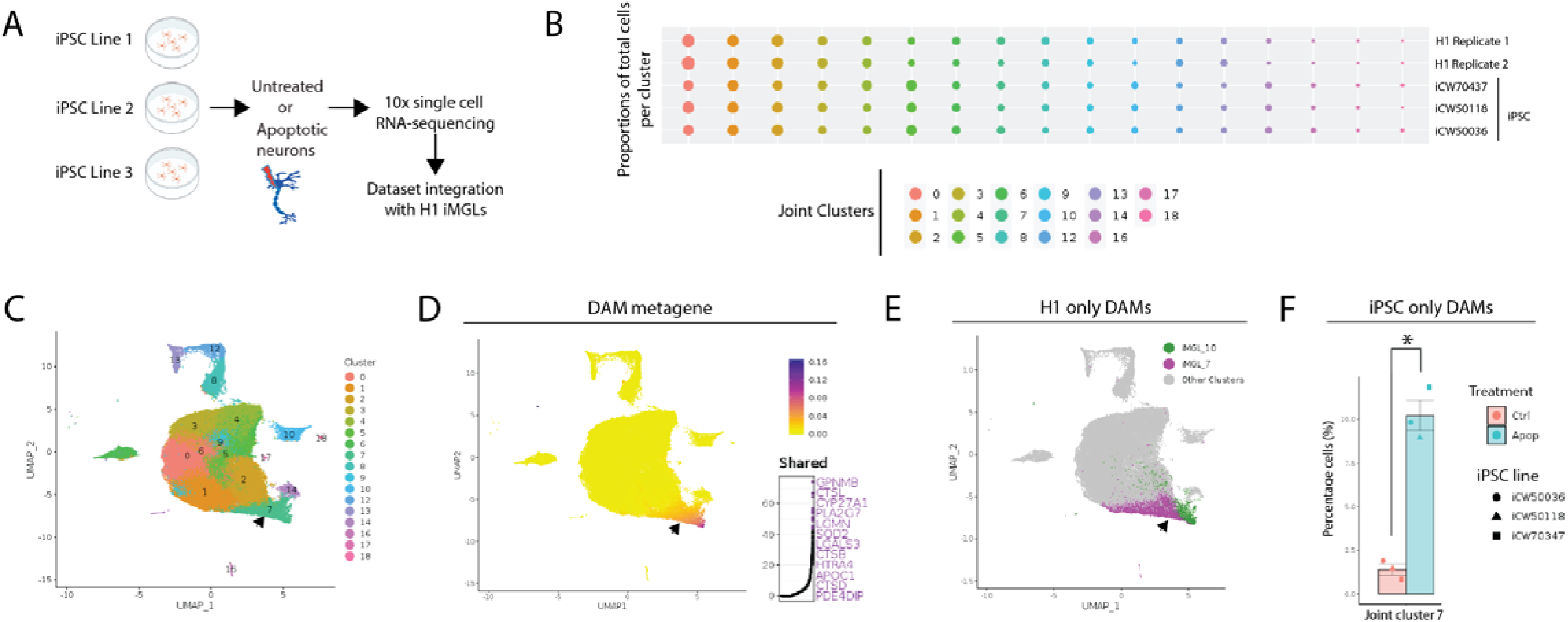
Multiple iPSC line-derived iMGLs also exhibit transcriptional diversity and DAM induction: A) Schematic of experiment, three separate iPSC lines were differentiated into iMGLs and either untreated or exposed to iMGLs followed by scRNAseq. Resulting data was integrated with H1 iMGL data from Figure 1. B) Proportion of cells per joint cluster for each dataset (iMGL H1 replicates or individual iPSC-derived iMGLs), dots are sized by proportion of total cells per dataset. C) UMAP projection of iMGL dataset integration, cells colored by cluster. D) Left, UMAP projection of LIGER integration showing the shared metagene common to both H1 and iPSC-derived iMGLs in cluster 7 of this integrative analysis. Right, top constituent genes of this shared factor. E) LIGER dataset integration, showing only H1 iMGLs with cluster iMGL_7 (magenta) and iMGL_10 (green) highlighted, both lie in join cluster 7 of this analysis. F) For iPSC-differentiated iMGLs, the percentage of cells in joint cluster 7 in control and apoptotic-neuron treated conditions (t-test, p-value = 0.00471). In C-E, black arrow highlights joint cluster 7.

## Discussion

As the principal macrophages cells of the brain parenchyma, microglia play essential roles in development, homeostasis and disease^1, 47^. These cells can form transcriptional subtypes, also known as states, in a context-dependent manner which presumably represent a diversification of function^26^ but the actual roles of these states are not known. Moreover, the recent identification of multiple disease variants mainly expressed in microglia highlighted the need for better characterization of these states specifically in humans. *In vitro* models can provide a powerful tool for functional dissection of many microglial genes and the impact of genetics risk due to their scalable and tractable nature but microglial cell lines and primary microglia do not accurately model their *in vivo* counterparts^15, 19^.

In this study, we overcame several technical obstacles to provide a toolbox for the study of human microglia states *in vitro*, as well as a single cell gene expression resource for the field that can be further validated and studied *in vivo* and in human chimeric and organoid models. Leveraging a new human microglia transcriptome dataset, we find that challenging iMGLs with CNS substrates produces a large degree of transcriptional diversity and strikingly, these *in vitro* states map to those found in the human brain. We find that exposure to apoptotic neurons or Aβ fibrils is sufficient to create a TREM2-dependent DAM-like state that is analogous to those found *in vivo* in human and mouse^7, 21, 48^. Therefore, addition of easily generatable substrates can produce models of human microglial states in a dish. Critically, we demonstrate that these features are shared over multiple iPSC lines, enabling investigators to analyze microglia from a full spectrum of patients and diseases and prioritize human cell lines for in depth functional characterization^46^.

An additional challenge of studying microglia is that, unlike other brain cell-types, these cells are not amenable to viral transduction or other forms of DNA delivery. The development of genetically modified iPSCs is low-throughput, costly and knock-out of essential genes can impact differentiation^49^. To further enable scalable experiments on iMGLs, we developed a lentiviral protocol that broadly transduces iMGLs in the last phase of their differentiation with modest perturbation of the core microglia transcriptional signature. We applied this technology to discover that MITF, a transcription factor upregulated in AD^23, 24, 45^, drives an increase in DAM genes and a phagocytic state when overexpressed in unstimulated iMGLs. This reveals a new function for MITF in regulating DAM gene signature and microglial function in neurodegenerative disease. By allowing for rapid and facile genetic modification of iMGLs, this technology will enable the identification of other key regulators of microglial states and functions^26, 31^, extensive characterization of the impact of human genetics (eg. targeting risk genes for AD) and the rapid application of molecular tools.

Although we focus on the DAM signature, the resources provided here will enable the dissection of many other microglial states such as interferon-responsive or proliferative microglia^6, 8, 10, 13, 17, 26, 29, 50^. By combining substrate exposure, scRNAseq, integrative analyses, epigenetics and viral vectors, it will be possible to scalably manipulate the genes or transcription factors that define these cell states and rapidly assay their impact on microglial functions. This platform also makes it possible to generate a large number of cells of each state to submit to unbiased characterization methods, such as proteomics or epigenetics, that are challenging to carry out on human tissue because of the small number of microglia that can be obtained from a single individual. Finally, combining substrate-exposed and/or virally modified iMGLs with human neuron and/or astrocyte co-cultures^51^ or with chimeric models^17, 33^ will enable the study of non-cell autonomous phenotypes and neuron-glial-immune interactions. In conclusion, our approach will enable a broad characterization of human microglia states and bridge the gap between transcriptomic and functional role of these states in neuro-immune interactions in health and disease.

## Supplementary Tables

Supplementary Table 1: Mean quality control (QC) values and number of cells per condition and replicate after QC

Supplementary Table 2: Differentially expressed genes per cluster for full iMGL dataset (H1)

Supplementary Table 3: Results of differential abundance testing of iMGL clusters across conditions. Adjusted p-values representing the differential abundance versus control for each substrate exposure for each cluster.

Supplementary Table 4: Shared gene signature in human and iMGL DAMs, hypergeometric test

Supplementary Table 5: Shared gene signature in human and iMGL DAMs, LIGER metagene

Supplementary Table 6: Differentially expressed genes between control and lentivirus treated iMGLs

Supplementary Table 7: iMGL ATAC-seq including DAR regions between iMGL AN and nontreated, list of transcription factors and motifs from HOMER analysis and overlap between HOMER and SCENIC analysis

Supplementary Table 8: Differentially expressed iMGL regulons identified by SCENIC

Supplementary Table 9: MITF bulk-RNA includes normalized gene expression data for all samples with statistics and fold change and overlap between MITF DEG and iMGL DAM

Supplementary Table 10: iPSC line information. Also includes scRNAseq summary data: Mean QC values and number of cells per condition per line after filtration for iPSC-differentiated iMGLs

## Data and code availability

All iMGL data will be deposited on GEO and Terra including raw and Cell Ranger output of iMGL (H1 and CW50118, CW500036 and CW70437) single cell RNA-sequencing, fastq and bam files of iMGL untreated and treated with apoptotic neurons for ATAC-seq and fastq and bam files of MITF overexpressing and mCherry control bulk RNA-sequencing. Any additional data and code is available from the corresponding authors.

## Online Methods

### Experimental Methods

#### Embryonic stem cell and iPSC lines

Except otherwise stated H1 embryonic stem cells (WiCell) were used. The following iPSC lines used in Figure 6 and Figure S12 were obtained from the CIRM hPSC Repository funded by the California Institute of Regenerative Medicine (CIRM): CW50118, CW50008, CW50065, CW500036 and CW70437. The cell lines chosen for scRNAseq (CW50118, CW500036 and CW70437) were confirmed as APOE 3/3 status with no TREM2 mutation by sanger sequencing. In addition, individuals did not exhibit cognitive decline at the time of collection which was after 70 years old for all lines (Supplementary Table 10), suggesting these lines should represent healthy individuals. The TREM2 knockout and isogenic control iPSC lines were previously characterized^21^ and obtained from Mathew Blurton-Jones. These lines were derived from cell line AICS-0036-006 from the NIGMS Human Genetic Cell Repository at the Coriell Institute for Medical Research thus have not undergone the standard quality control of the Repository.

#### iMGL differentiation

iMGLs were differentiated as previously described^22^. Briefly, iPSC or ESC were cultured in Essential 8 (E8) (Thermo Fisher Scientific) media on Matrigel (Corning) coated 6-well plates. When confluent, cells were dissociated using Accutase (Stem Cell technologies), centrifuged for 5mins at 300xg and counted using trypan blue (Thermo Fisher Scientific). Next, 200,000 cells/well were resuspended in E8 containing 10uM Y27632 ROCK inhibitor (Selleckchem) in low adherence 6-well plate (Corning). For the first 10 days, cells were cultured in HPC medium [50% IMDM Thermo Fisher Scientific), 50% F12 (Thermo Fisher Scientific), ITSG-X 2% v/v (Thermo Fisher Scientific), L-ascorbic acid 2-Phosphate (64 ug/ml, Sigma), monothioglycerol (400mM, Sigma), Poly(vinyl) alcohol (PVA) (10mg/ml, Sigma), Glutamax (1X, Thermo Fisher Scientific), chemically-defined lipid concentrate (1X, Thermo Fisher Scientific) and non-essential amino acids (Thermo Fisher Scientific)]. At day 0, embryoid bodies (EB) were gently collected, centrifuged at 100xg and resuspended in HPC medium supplemented with 1uM ROCK inhibitor, FGF2 (50 ng/ml, Thermo Fisher Scientific), BMP4 (50ng/ml, Thermo Fisher Scientific), Activin-A (12.5ng/ml, Thermo Fisher Scientific) and LiCL (2 mM, Sigma), then incubated at in hypoxic incubator (5% O2, 5% CO2, 37°C). On day 2, cells were gently collected and the media changed to HPC medium supplemented with FGF2 (50ng/ml, Thermo Fisher Scientific) and VEGF (50 ng/ml, PeproTech) and returned to the hypoxic incubator. On day 4 cells were gently collected and media changed to HPC medium supplemented with FGF2 (50ng/ml, Thermo Fisher Scientific), VEGF (50 ng/ml, PeproTech), TPO (50 ng/ml, PeproTech), SCF (10ng/ml, Thermo Fisher Scientific), IL6 (50ng/ml, PeproTech) and IL3 (10ng/ml, PeproTech) and incubated in normoxic incubator (20% O2, 5% CO2, 37°C). At day 6 and 8, 1 ml of day 4 media was added in each well. On day 10, cells were collected, counted using trypan blue and frozen in Cryostor (Sigma Aldrich) in aliquots of 300,000-500,000 cells.

To start iMGL differentiation, cells were thawed, washed 1x with PBS and plated at 100,000-200,000 cells per well in 6-well plate coated with matrigel in iMGL media [(DMEM/F12 (Thermo Fisher Scientific), ITS-G (2% v/v, Thermo Fisher Scientific), B27 (2% v/v, Thermo Fisher Scientific), N2 (0.5% v/v, Thermo Fisher Scientific), monothioglycerol (200 mM, Sigma), Glutamax (1X, Thermo Fisher Scientific), non-essential amino acids (1X, Thermo Fisher Scientific)] supplemented with M-CSF (25 ng/ml, PeproTech), IL-34 (10 ng/ml, PeproTech) and TGFB-1 (50ng/ml, PeproTech). Cells were fed every 2 days and replated at day 22. On day 30, cells were collected and replated in iMGL media supplemented with M-CSF (25 ng/ml, PeproTech), IL-34 (10 ng/ml, PeproTech), TGFB-1 (50ng/ml, PeproTech), CD200 (100ng/ml, VWR) and CX3CL1 (100ng/ml, PeproTech). Cells were used at day 40 for functional and transcriptomic assays. iMGL differentiation was assessed at day 10 (expression of CD43 and CD45) and day 40 (expression CD45, CD11B, P2RY12 and Cx3CR1) by flow cytometry.

#### Flow cytometry

For sample processing, iMGLs were detached using cold PBS, then resuspended in FACS buffer (PBS containing 2%BSA and 0.05mM EDTA). Samples were incubated for 15 mins in human Fc block (BD Biosciences) followed by 1h staining with conjugated antibodies (see below) at 4C. Samples were washed 3x with FACS buffer and resuspended in 500ul of FACS buffer for flow cytometry. Samples were run on CytoFLEX S analyzer (Beckman Coulter) until at least 2,000 cells were recorded. For analysis, cells were identified according to the following gating: 1) Cell vs debris (FSC-A vs SSC-A), 2) singlets (FSC-A vs FSC-H), and antibody specific gating based on negative control sample. Antibodies for staining iMGLs were CD45-FITC (BioLegend), CD11B-APC-750 (BioLegend), P2RY12-PB450 (BioLegend), Cx3CR1-PrCp (BioLegend) 1/500 for all.

For phagocytosis, cells were treated with pHrodo labeled substrates (see below) for 24h. iMGL were then washed with PBS and collected in FACS buffer (PBS containing 2%BSA and 0.05mM EDTA). Samples were then directly run on CytoFLEX S analyzer (Beckman Coulter) until at least 2,000 cells were recorded. For analysis, cells were identified according to the following gating: 1) Cell vs debris (FSC-A vs SSC-A), 2) singlets (FSC-A vs FSC-H), 3) viability (based on DAPI stain). Phagocytosis was quantified as mean fluorescent intensity of pHrodo of each sample. Each experiment was done on multiple independent differentiations and each dot in the graph represents a biological replicate.

All analyses were performed using FlowJo V10 and statistical analysis performed using Prism9.

#### CNS substrate isolation and iMGL treatment

Note: Unless otherwise stated, treatment of iMGLs with CNS substrates was for 24 hours prior to transcriptomic, flow cytometry or RNAscope analysis.

Synaptosomes were prepared from rodent brains as described previously^52^. Briefly, C57BL/6J mice were euthanized with CO2, then brains were dissected and homogenized in HEPES-buffered sucrose (0.32 M, 5 mM HEPES, pH 7.4). The resulting homogenate was spun at 800-1200xg to separate the nuclear fraction. A further spin at 15,000xg was carried out to generate crude synaptosomes used for future experiments.

Myelin was isolated from C57BL/6J mice. Animals were transcardially perfused with cold HBSS and whole brains were manually homogenized in RPMI. Samples were applied to a percoll gradient, and after a 30 min spin at 500xg, the top layer was collected. Myelin was washed twice with water and used for future experiments.

Apoptotic neurons were generated using either SH-SY5Y cells or iNeurons generated as described^48^. Using neurogenin-2 (NGN2) for 14 days. Briefly, stem cells were grown in StemFlex (Stem Cell Technologies) and grown on Matrigel (Corning) coated plates at 37°C and 5% CO2. Cells were infected with TetO-Ngn2-Puro, and rtTA lentiviral particles in StemFlex medium with 1 µM ROCK inhibitor Y-27632 for 24 hours, were then passaged and differentiation was started when cells reached 70-80% confluency. For the first 2 days, cells were grown in N2 media (DMEM/F12 (Life Technologies, 11320-033), N2 supplement (0.5%v/v, Gibco), 1X GlutaMAX (Gibco), 0.1mM Non-essential amino acid (Gibco), 0,5% glucose, doxycycline hyclate (2 µg/mL)). Day 0, N2 media was supplemented with 0.1uM LDN, 5uM XAV, 10uM SB, and on day 2, the cells were fed with N2 media, no supplement. At day 3, cells were replated to 100 000 cells/ well in 6-well plate and transferred to Neurobasal media (Neurobasal (Gibco), 1X B27 (Gibco), 1X GlutaMAX (Gibco), 0.1mM Non-essential amino acid (Gibco), 0,5% glucose, doxycycline hyclate (2 µg/mL)) supplemented with 10ng/ml CNTF, 10ng/ml BDNF and 10ng/ml GDNF. Cells were fed every other day until day 14. Mature cells were submitted to UV radiation (500 J/m2 using a UV Crosslinker) and gently collected 24h later and used for future experiments.

For conjugation to pHrodo (Red SE or Green, STP Ester; Thermo Fisher Scientific), labeling was done according to manufacturer’s protocol. Briefly, substrates were incubated with pHrodo for 2 hr at RT protected from light at 1 mL pHrodo/1mg substrate, washed 10x using PBS and frozen in 5% DMSO. Only synaptosomes, myelin or apoptotic neurons were conjugated.

Amyloid fibrils were prepared from fluorescently-labeled (Beta-Amyloid (1-42) HiLyte Fluo, AS-60479-01, Anaspec). Aβ peptides corresponding to amino acids 1–42 were dissolved in sterile water to 100ug/ml concentration, vortexed thoroughly and shaken in a 37°C incubator for 4 hours. Then the Aβ peptides were incubated for 5 days at 37°C to fibrilize. Amyloid fibrils were then mixed and delivered to cells at 5 μg/mL. Fresh amyloid fibrils were created for each experiment.

#### Sample preparation for single cell RNA sequencing

For single cell sequencing experiments, iMGLs at day 40 were treated with myelin, synaptosome, apoptotic neurons or amyloid fibrils for 24h (see above). Cells were washed with warm media to remove floating cells, and detached in PBS on ice. Cells were then centrifuged and resuspended in cold PBS at 1000 cell/ul.

For the single cell sequencing experiment comparing control versus lentivirus-transduced cells, iMGLs were transduced with a non-targeting (NT) gRNA-Cas9-mCherry lentivirus (see below for more details on lentiviral vector, production and exposure) on day 35. Cells were exposed to the virus overnight, and 100% of the medium was changed the next morning. Cells were then cultured for 7 more days and fed on a regular schedule. On the 7th day, mCherry expression was confirmed, and all media was removed. Cells were washed with warm media to remove floating cells, and detached in PBS on ice. Cells were then centrifuged and resuspended in PBS.

After resuspension and counting, cells loaded into the 10x Chromium V3 system (10x Genomics). Reverse transcription and library generation were performed according to the manufacturer’s protocol. Due to a counting error, the number of cells loaded for replicate 1 of the H1 scRNAseq dataset were ∼3-fold lower than the replicate 2, necessitating a data integration strategy for batch correction (see below). Sequencing was performed on a Novaseq S2 (Illumina).

#### Real-time quantitative PCR

iMGLs day 40 were treated for 24h and collected using RLT buffer. RNA extraction was done using RNease Plus mini kit (Qiagen) and done according to manufacturer’s protocol. For quantitative RT-PCR, TaqMAn RNA-to-Ct 1-step kit (Thermo Fisher Scientific) was used according to manufacturer’s protocol using the following Taqman probes (Thermo Fisher Scientific): GAPDH (HS02786624_G1) and GPNMB (HS01095669_M1). Quantification was done using 2 ^- ΔΔCT^ method^53^. Each biological sample was measured in triplicates and the average of each biological sample is shown in graph. Statistics were done using Prism9 software and graph are showing individual cells with mean and standard deviation. Two-tailed t-test was used to determine significance.

#### Immunocytochemistry, imaging and quantification

iMGL plated on matrigel coated coverslips were fixed in 4% PFA for 15 mins, followed by permeabilization (0.2% triton-X in PBS) for 10- mins and blocking (5% Normal Donkey serum in 0.025% Triton-X/ PBS) for 30 mins. iMGLs were incubated with primary antibodies overnight (concentrations below) and washed 3x for 10 mins with 0.025% Triton-X/PBS. Then cells were incubated with a secondary antibody (Thermo Fisher Scientific) for 60 mins at room temperature. Following 3x washes for 10 mins each with 0.025% Triton-X/PBS, coverslips were mounted on slides using ProLong Gold antifade (Thermo Fisher Scientific).

Primary antibodies used and concentration: TREM2 (R&D systems, AF1828-SP) 1/100, GPNMB (Cell Signaling, E1YZJ) 1/500

Secondary antibodies used and concentration: Donkey anti-Rabbit 488 (Thermo Fisher Scientific) 1/500, Goat anti-Rabbit 594 (Thermo Fisher Scientific) 1/500

Imaging was performed on an Andor CSU-X spinning disk confocal system coupled to a Nikon Eclipse Ti microscope equipped with an Andor iKon-M camera. Images were acquired using a 60x oil objective (Nikon). All images shown are representative images taken from at least 3 independent experiments. Statistics were done using Prism9 software and graphs showing individual cells with mean and standard deviation. Two-tailed t-test was used to determine significance.

#### In situ hybridization RNAscope and quantification

RNAscope (Advanced Cell Diagnostics) was carried out according to the manufacturer’s protocol for cultured adherent cells on coverslips. Briefly, iMGL plated on matrigel coated coverslips were fixed for 30 mins using 4% PFA and dehydrated (5 min 50% ETOH, 5min 70% ETOH, 2x 5min 100% ETOH) and kept at -20C until use. When ready to treat, cells were underwent the RNAscope Multiplex Fluorescent V2 Assay (Advanced Cell Diagnostics) and probes against human APOE (cat #433091), ABCA1 (cat # 432291) and C1QA (cat # 485451-C2) were used at recommended concentrations. TSA Plus fluorophores (Perkin Elmer) were used at 1:1500 concentration.

Imaging was performed on an Andor CSU-X spinning disk confocal system coupled to a Nikon Eclipse Ti microscope equipped with an Andor iKon-M camera. Images were acquired using a 60x oil objective (Nikon). C1QA was used to identify regions of interest (ROI) to identify individual cells. Because microglia change shape and can become ameboid in response to stimulus, quantification of intensity could not be normalized to the cell’s area, therefore, only fluorescent intensity of target probe was measured using FIJI^54^. Statistics were done using Prism9 software and graphs showing individual cells with mean and standard deviation. Two-tailed t-test was used to determine significance.

#### Production of lentivirus and Vpx VLPs

Lentivirus and Vpx VLPs were produced by transfecting HEK293T cells using TransIT-LT1 reagent (Mirus). For lentivirus, the following plasmids were transfected (amounts per well of a 6-well plate): 1.6 μg mCherry- and Cas9-expressing lentiviral genome plasmid pXPR_BRD044 (obtained from the Broad Institute Genome Perturbation Platform), 0.4 μg pCMV-VSV-G (Addgene #8454) and 1 μg psPAX2 (Addgene #12260). For Vpx VLPs 0.4 μg pCMV-VSV-G and 2.6 μg pSIV3 Vpx plasmids were transfected. Media was changed to OptiMEM 18 hours after transfection (2 mL per well in a 6-well plate). Two days after transfection, viral supernatant was harvested and centrifuged at 500xg for 10 min to remove any cells in the supernatant. Clarified supernatant was concentrated 10X by incubating with Lenti-X Concentrator (Takara) overnight (due to the size of the viral genome the virus had to be concentrated to achieve sufficient titer). The mixture was spun down the next day per manufacturer’s instructions and resuspended in base iMGL media. Vpx VLPs were not concentrated. Lentivirus was titrated using Lenti-X p24 Rapid Titer Kit (Takara). Both lentivirus and Vpx VLPs were flash frozen and stored at -80°C before use.

#### Lentivirus transduction for FACS, single cell sequencing and RT-qPCR time-course studying the effects of lentiviral transduction of iMGLs

For FACS, iMGLs were seeded in a 96-well plate. On day 35 lentivirus and Vpx VLPs were added. The next day, all media was removed and replaced with fresh media and then changed every 2 days. On day 42 cells were detached as described above and FACS was performed to quantify mCherry expression from lentivirus. For single cell sequencing, iMGLs were seeded in a 6-well plate. One 6-well plate well corresponds to one replicate for single cell sequencing. 200,000 cells were seeded and treated with lentivirus and Vpx VLPs as described above. Controls were treated the same as lentivirus in terms of media changes. 7 days after virus addition, cells were detached with cold PBS on ice for 10 minutes and spun at 300xg for 5 minutes. Cells were then resuspended in a small volume of PBS and counted. Approximately 20,000 cells were used as input for each single cell sequencing replicate.

For the time course studying expression of interferon-stimulated genes after exposure to lentivirus, iMGLs were seeded into 24-well plates. On day 35 cells were co-transduced with lentivirus and Vpx VLPs. RNA was harvested on days 37, 42 or 42. To harvest RNA, iMGL media was removed and RLT Plus buffer from RNeasy kit (Qiagen) was added directly to the wells. RNA was purified using manufacturer’s instructions. For quantitative RT-PCR, TaqMan RNA-to-Ct 1-step kit (Thermo Fisher Scientific) was used according to manufacturer’s protocol using the following Taqman probes (Thermo Fisher Scientific): GAPDH (HS02786624_G1), IFI6 (Hs00242571_m1), IFITM3 (Hs03057129_s1) and ISG15 (Hs01921425_s1). Data was analyzed using CFX Maestro software (BioRad).

#### ATAC-seq library preparation and sequencing

ATAC-seq libraries were prepared essentially as described, with the following modifications. For each sample, 10,000 viable frozen cells were used per library construction. We optimized the number of PCR cycles by stopping PCR after 5 cycles, taking an aliquot from the partially amplified library, and performing qPCR for 25 cycles. We visually examined the qPCR amplification profile to determine the number of cycles required to reach 1/3 of the plateau, and extend the original PCR by this number of additional cycles. Libraries were sequenced on an Illumina Next-seq platform using a 75 cycle Nextseq 500 high output V2 kit (Read 1: 36 cycles, Index 1: 8 cycles, Index 2: 8 cycles, Read 2: 36 cycles).

#### MITF overexpression lentivirus

For MITF overexpression, lentiviral vectors contained either MITF (NM_198159.3) or mCherry under the EF1a promoter. Lentivirus was produced as described above. At day 35, iMGLs were infected with MITF mCherry expressing lentivirus and the media was changed the following day. At day 42, iMGLs were treated for functional or transcriptomic characterization.

#### Bulk RNA-seq library preparation and sequencing

iMGLs were collected using RLT buffer (Qiagen) and RNA extraction was done using the RNeasy Plus mini kit (Qiagen) and done according to the manufacturer’s protocol. Quantification and RNA integrity was assessed using the RNA 600 Pico chip on a Bioanalyzer (Agilent). 5ng of RNA was used as input for library constructions. Libraries were constructed using NEBNExt Poly(A) mRNA Magnetic isolation module (NEB E7490) in combination with NEBNext Ultra RNA library prep for Illumina (NEB E7530) following the manufacturer’s instructions. Libraries were quantified using Agilent high sensitivity DNA chip and sequenced on NextSeq (Illumina). Samples were sequenced in 4 flow chambers and reads from each were merged to generate a final fastq final for each sample.

### Analysis Methods

#### Single Cell RNAseq Preprocessing, quality control and general analysis

Cellranger (version 3.1, 10X Genomics) was used for demultiplexing, barcode preprocessing, generation of fastq files, alignment (to GRCh38-2020-A/GENCODE v32/Ensembl 98) and counting of unique molecular identifiers (UMI). Due to differences in sequencing depth between replicates (see above), we applied Cellranger’s aggr function to all samples to merge and normalize by sequencing depth.

All downstream analysis (see below for specific analyses) was performed in R using the LIGER (version 0.5) and Seurat (either version 2.3.4 or 3.2.1) packages^28, 55^. Note that lentivirus-exposed iMGLs were excluded from this analysis and analysis of this experiment is described below. All data and metadata (replicate and condition) were incorporated into a single object and the percentage of mitochondrial RNA per cell was calculated. We only considered cells with the following characteristics: 1) Number of Genes: 2000-7000, 2) Number of UMIs: 500-6000, 3) Percentage mitochondrial RNA 0-0.2%. The total number of cells per condition after quality control is reported in Supplementary Table 1.

Cells of interest were subsetted and converted to a LIGER object for integration. To integrate the two replicates, we considered the replicate (1 or 2) as the LIGER dataset variable and applied the LIGER pipeline of dataset integration (k=20, lambda= 5). Clustering was performed using the louvainCluster function (clustering resolution=0.7). A UMAP^56^ embedding was calculated using LIGER’s runUMAP function. The alignment was evaluated with the CalcAlignment and CalcAgreement LIGER functions. All gene expression, metagene and violin plots were produced using LIGER functions and modified using ggplot2.

#### Differential expression analysis

To identify differentially expressed genes per LIGER cluster we used the MAST package^57^ and included the number of UMIs and number of genes as covariates. Only genes expressed in more than 10% of cells per cluster were considered. Cell gene expression was then scaled, centered and averaged per cluster prior to heatmap generation. Heatmaps were generated using the ComplexHeatmap R package^58^.

#### Module score for artifactual activation of microglia (ExAMs)

The module score for testing for artifactual gene expression was created using genes previously identified as being upregulated during single-cell isolation and cell handling^27^. The genes were: *RGS1*, *HIST2H2AA1*, *HIST1H4I*, *NFKBIZ*, *KLF2*, *JUNB*, *DUSP1*, *CCL3*, *HSPA1A*, *HSP90AA1*, *FOS, HSPA1B*, *JUN*, *JUND*, *NFKBID, GEM*, *CCL4*, *IER5*, *TXNIP*, *HIST1H2BC*, *ZFP36*, *HIST1H1C*, *EGR1*, *ATF3*, *RHOB*. The module score was implemented in Seurat using the AddModuleScore function with a control size of 25.

#### Cluster proportions, differential abundance test and fold change analysis

For plotting cell proportions the percentage contribution of each condition to a cluster by replicate was calculated and averaged. For determining statistical significance of cluster abundances across conditions, we used the counts (not proportions) and implemented a Dirichlet regression, a multivariate test which accounts for overall composition per sample. This was implemented with the DirichletReg R package (https://github.com/maiermarco/DirichletReg) and a previously published workflow ^59^. Fold change values were calculated by replicate, relative to the control condition and a log2 transformation for final log fold change plots reported.

#### Gene expression per exposure condition and GO analysis

For GO analysis per exposure condition, the MAST package^57^ was applied to the integrated Seurat object (see above), using the exposure condition as a variable. For Gene Ontology (GO) analysis, genes with a log fold change >0 and a final adjusted p-value <0.05 were used. The compareCluster function from clusterProfiler package^60^ used to perform and plot differential GO pathway analysis.

#### LIGER integration of iMGLs with different datasets

For large dataset integrations (ie. with large numbers of cells, individuals or both) LIGER for scalability. Analysis of the iNPH, the final microglia object and cluster labels from Kamath et al. (In preparation) were used. For analysis of the mouse DAM dataset, the data was reprocessed and analyzed using Seurat (3.2.1) and cluster identities (in particular the identification of DAM microglia) was determined based on differential gene expression and comparison with source paper (eg. upregulation of *Lpl*, *Clec7a*, *Trem2*, *Itgax*, *Cd9*, *Axl*)^7, 33^. For xMG analysis, the cluster assignments from the original paper were used ^33^.

All cross-context (xenotransplanted iMGLs) and cross-species (human in-vivo and mouse in-vivo) dataset integrations were performed using the LIGER package^28^. Post-QC iMGLs count data was used for all alignments. For each alignment, the data, parameters and dataset variable arrangement was as follows:

| <u>Data</u> | <u>Reference</u> | <u>Parameters</u> | <u>Dataset variable</u> |
| --- | --- | --- | --- |
| Human *in vivo*: iNPH | Kamath et al., in Prep | k=20, lambda=5, clustering resolution=0.7 | iMGL replicates (untreated and all CNS substrates, individual patients (58 datasets)) |
| Human *in vivo*: iNPH | Kamath et al., in Prep | k=20, lambda=5, clustering resolution=0.7 | iMGL replicates (just untreated iMGLs, individual patients (58 datasets)) |
| Mouse *in vivo*: DAMs | Keren-Shaul et al., 2017 | k=20, lambda=5, clustering resolution=0.5 | iMGL replicates, mouse microglia as one dataset |
| Xenotransplanted iMGLs: xMGs | Hasselmann et al., 2019 | k=20, lambda=5, clustering resolution=0.6 | iMGL replicates, xMG in WT, xMG in 5xFAD |

Note: LIGER alignment of H1 iMGLs with iPSC-derived iMGLs (Figure 6) is described below.

#### GSEA analysis of iMGL differentially expressed genes and other datasets

The GSEA analysis^61^ was run using the fgsea package^62^ on differentially expressed genes identified above (see: Differential expression analysis). Differentially expressed genes for each iMGL cluster were ordered by log (fold change). Pathways were defined as positively differentially enriched genes identified by differential expression analysis from other datasets: Gerrits et al. 2021, Kamath et al. In preparation. Data was plotted using the ComplexHeatmap R package^58^.

#### Hypergeometric test comparing human and iMGL DAM gene expression

Prior to performing the hypergeometric test, DAM genes in both iMGLs and *in vivo* human were identified. DAM genes in iMGLs were determined by taking the union of positively enriched, differentially expressed genes for iMGL_07 and iMGL_10 (see “<u>Differential expression analysis”</u> above). To determine the number of differentially expressed DAM genes in the in-vivo dataset, we performed differential expression analysis using MAST^57^ of human microglia in the DAM cluster versus all others. We included the number of UMIs and number of genes as covariates. The hypergeometric test was run on intersecting genes using the dhyper function from the Stats R package. The background number of genes for each dataset was calculated based on genes expressed in more than 1% of cells (13,187 genes).

#### Cluster occupancy test

To test the statistical significance of the co-clustering of iMGL_7, iMGL_10 and human DAMs (Microglia_GPNMB_LPL), a binomial test was applied to determine if cells from each group were enriched in joint clusters 8 and 15 relative to other clusters. The binom.test function from Stats R package was used for this analysis.

#### Monocle trajectory analysis

For trajectory analysis of apoptotic neuron and amyloid beta-exposed iMGLs, normalized gene expression data was aligned by replicate using Batchelor^63^, which was chosen due to compatibility with the Monocle3 pipeline. Dataset integration across replicates with Batchelor gave similar results as LIGER and Seurat (data not shown). The preprocessing, dimensionality reduction and pseudotime ordering of cells was performed using Monocle3^34^ and cluster labels from initial LIGER analysis (see above) were used for analysis and plotting.

#### SCENIC transcription factor analysis

The post-QC, normalized iMGL gene expression matrix was used as a starting point for analysis with SCENIC^44^. Cells from the same LIGER cluster were randomly averaged into pseudocells of 20, which was previously shown to yield more robust results^64^. GENIE3 was used to identity TF/gene coexpression networks, followed by regulon analysis (runSCENIC_2_createRegulons) and regulon scoring per cell (runSCENIC_3_scoreCells) to create a regulon expression matrix for all cells. For the final list of significant regulons, a differential expression test was performed with the presto R package (wilcoxauc function) (https://github.com/immunogenomics/presto) and results were filtered by an adjusted p-value<0.01 and area under the curve (AUC)>0.7. Regulons were scaled and centered for plotting using the pheatmap package (https://github.com/raivokolde/pheatmap).

#### Lentivirus-exposed versus control iMGLs

Sequencing data was processed with Cell Ranger 3.1.0 (10x Genomics). Fastq files were mapped to a modified reference genome (GRCh38-2020-A/GENCODE v32/Ensembl 98) containing the mCherry gene. Count data was obtained using Cellranger count function. The count data was further analyzed using Seurat 3.2.2^55^. Count matrices from four conditions were merged, filtered (only cells with 2-7,000 genes, 500-60,000 reads and less than 20% mitochondrial RNA were kept) and normalized using standard log-normalization. Integration was performed using FindIntegrationAnchors and IntegrateData functions. To analyze the number of cycling cells, the CellCycleScoring function was used. To analyze the remaining gene expression changes in the dataset, cycling cells were removed (only cells in G1 phase were kept), the data was aggregated and differentially expressed genes were identified using Deseq2 1.30.1 package^65^ using default parameters (except for FDR = 0.01). Enriched gene ontology terms were annotated with the enrichR package using database GO_Biological_Process_2015^66^.

#### Analysis of iPSC-derived iMGLs and dataset integration

For analysis of lines CW50118, CW500036 and CW70437, single cell gene expression matrices were first filtered for low quality cells. As above, we only considered cells with the following characteristics: 1) Number of Genes: 2000-7000, 2) Number of UMIs: 500-6000, 3) Percentage mitochondrial RNA 0-0.2%. After this cell filtration step, cells were merged with the post-QC CNS-substrate exposed H1 iMGL scRNAseq dataset above (Figure 1). To integrate the two replicates, we considered the H1 iMGL replicate (1 or 2) and the iPSC stem cell line as the LIGER dataset variable and applied the LIGER pipeline of dataset integration (k=20, lambda=6). Clustering was performed using the louvainCluster function (clustering resolution=0.5). A UMAP^56^ embedding was calculated using LIGER’s runUMAP function. The alignment was evaluated with the CalcAlignment LIGER functions. Two small clusters (15 and 11) were enriched for neuronal genes and specific to the apoptotic neuron-treated conditions for all lines. These were considered doublets and removed prior to visualization and downstream analyses. All gene expression, metagene and violin plots were produced using LIGER functions and modified using ggplot2.

#### Bulk RNA-seq analysis

Reads were aligned to the human genome (hg19) using Picard (https://broadinstitute.github.io/picard/). Read counts were obtained using Featurecounts^67^, ComBat-seq was used to adjust for batch effects^68^ and DESeq2 used for differential expression^65^. For the hypergeometric test, the background number of genes for each dataset was calculated based on genes expressed in more than 1% of cells (13,187 genes).

#### ATAC-seq data processing

Reads were aligned to the human reference genome (hg19) with the BWA aligner^69^. Duplicates were removed using Picard tools (MarkDuplicates) (https://broadinstitute.github.io/picard/) and peaks called using HOMER 4.11.143 in “histone” mode. Diffbind^70^ was used to identify differentially represented regions between conditions.

#### Graphic design

Parts of several diagrams were created using <u>BioRender.com</u>.

## Acknowledgments

We thank Ben Deverman, Borislav Dejanovic and all members of Stevens and Macosko labs for helpful discussion when preparing the manuscript. We also thank the members of the International Neuroimmune Consortium for their insightful conversations and support in this project. This work was supported in part by: Alzheimer’s Association (ADSF-21-831114-C)(BS); Stanley Center for Psychiatric Research (BS); Broad Institute (V2F)(BS); Cure AD fund (CAF)(BS, MBJ and CKG); OneMind/Alzheimer’s Foundation of America (BS); HHMI (BS). M-JD is an Open Philanthropy Project Awardee of the Life Sciences Research Foundation. MT is recipient of FRQS and CIHR postdoctoral fellowships. TK is supported by the F30 (NIH, national institutes of aging) F30AG069446-01. VL is funded by Kuopio University Hospital VTR Fund, Academy of Finland (grant number 339767) and Sigrid Juselius Foundation. Isogenic TREM2 knockout iPSC lines were generated by the UCI-ADRC iPS cell core supported by NIH P30-AG066519. Experiments using the GFP-expressing iPSC line AICS-0036 were made possible through the Allen Cell Collection, available from Coriell Institute for Medical Research.

## Contributions

study design: MT, SJ, M-JD and BS

performed experiments: MT, SJ, M-JD with help from TA, NL, SM, CMW, CBE

performed analysis: M-JD, SJ and MT. Assistance provided by TK, SEM, AG and EZM

wrote the manuscript with input from all co-authors: M-JD, MT and SJ

provided reagents: JJ, FL, FZ, MBJ, BL

provided human tissue and snRNAseq data: VL, TK and EZM advised on methodology: NH, MBJ, BL, KE, CBE, BEB, CKG, FZ

project supervision: EZM and BS

---

**Figure S1:**
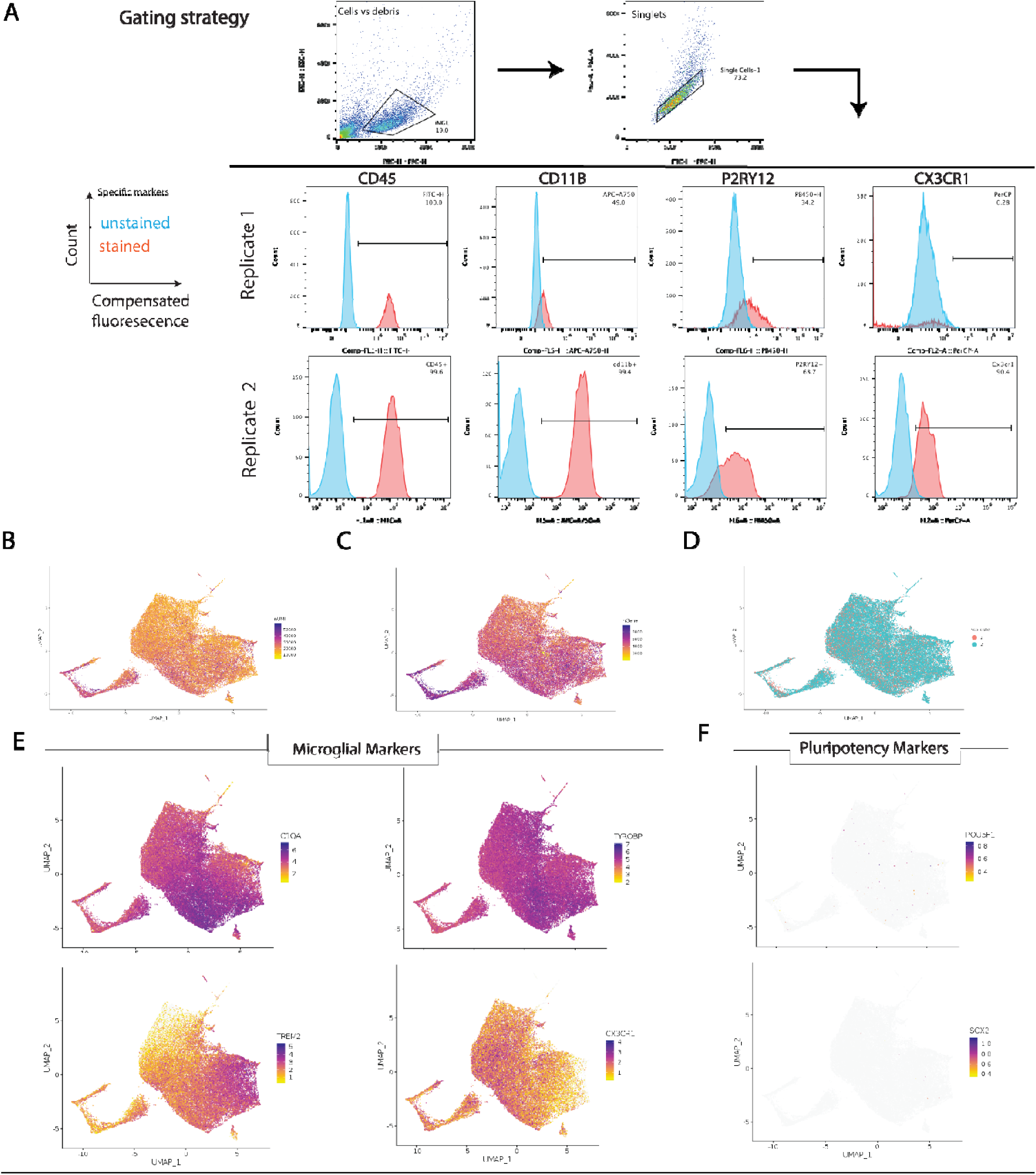
Single-cell RNAseq of iMGLs: Quality control. A) Flow cytometry gating strategy and plots illustrating expression of microglial protein markers from the two differenciations used for scRNAseq dataset. B-C) UMAP projection of quality control data for scRNAseq highlighting the number of unique molecular identifiers (nUMI, B) and the number of genes (nGene, C) for all cells showing reads and transcriptional diversity. D) UMAP projection of single iMGL transcriptomes plotted by replicate. E) UMAP projection of iMGLs showing expression of four key microglial markers. F) Lack of expression of pluripotency genes at single cell level. n=2 independent differentiations for B-F.

**Figure S2:**
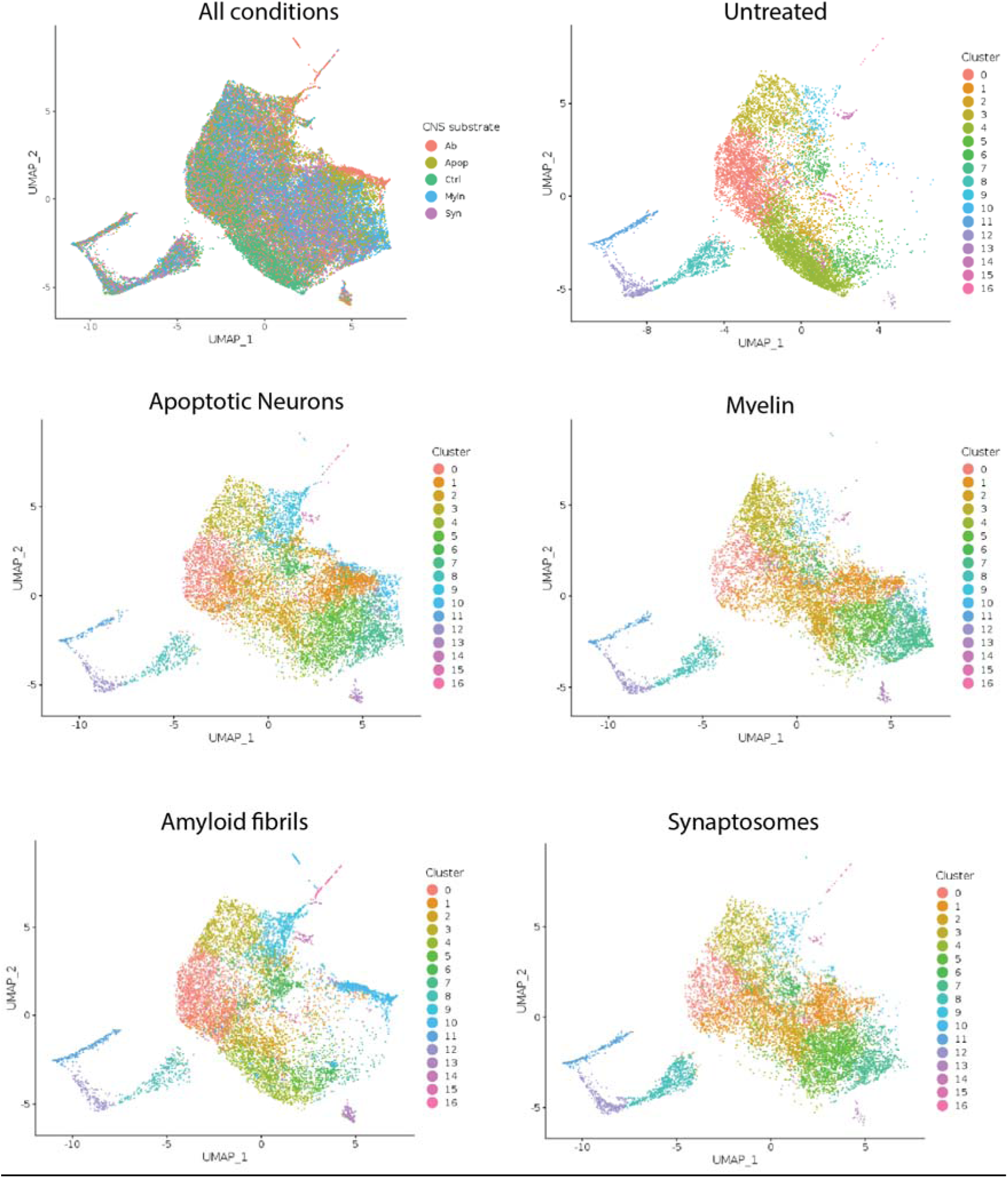
Single-cell RNAseq data for each condition. UMAP projection of single iMGLs for each condition (n=2 independent differentiations per condition), cells are colored by conditions (Top left), or clustering from figure Figure 1B.

**Figure S3:**
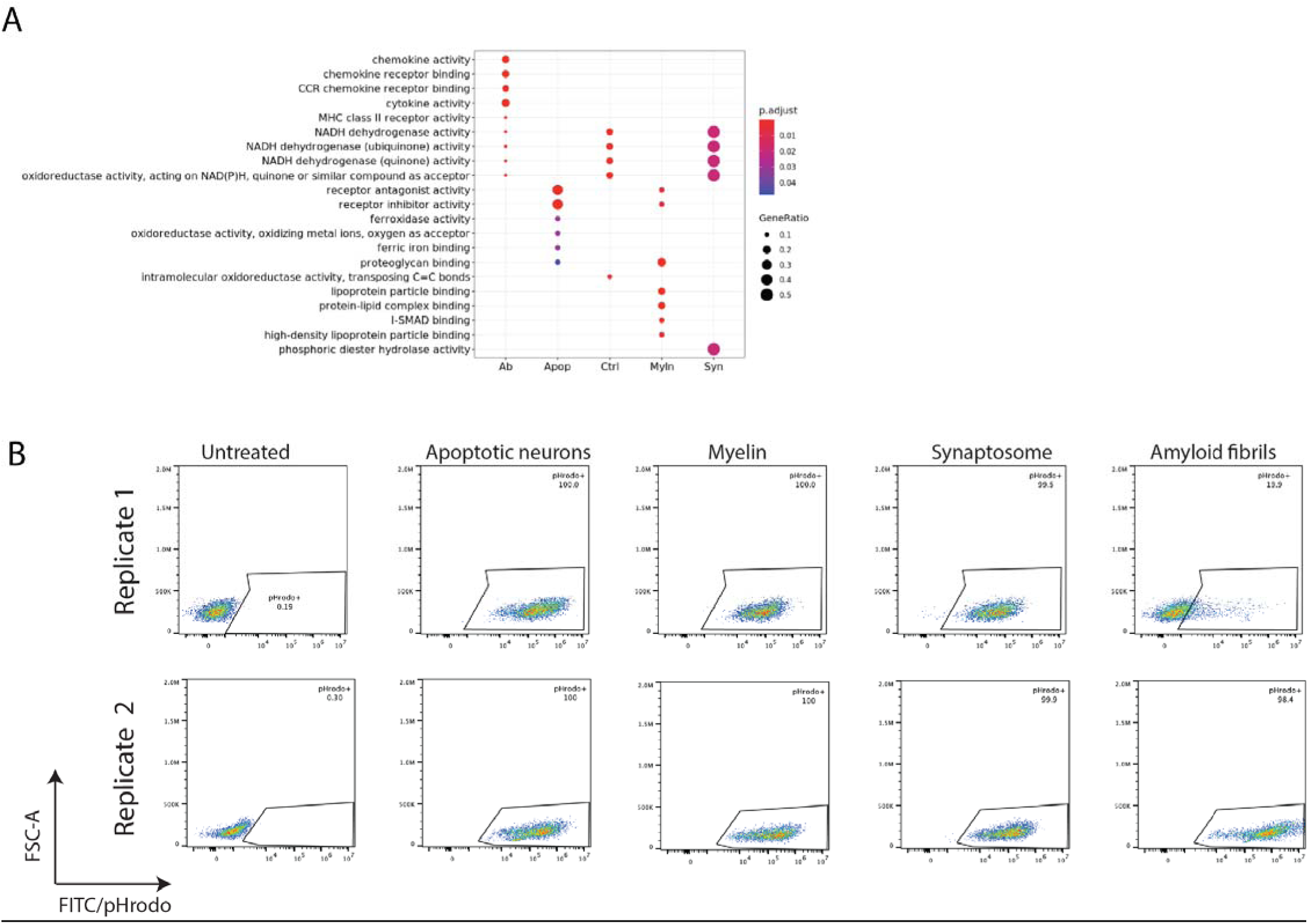
Transcriptional states are not explained by phagocytosis of substrates. A) Gene Ontology analysis of pseudobulk gene expression data (generated from scRNAseq data, see methods) plotted per condition reveals changes in functional pathways i duced by different substrates. (abbreviation Ab= synthetic A fibrils, Apop = Apoptotic neurons, Myln= Myelin, Syn= Synaptosome) B) Phagocytosis performed on iMGLs exposed to pHrodo-conjugated substrates (Apoptotic neurons, myelin, synaptosome) or 488 fluorescently-conjugated Amyloid-beta reveals complete phagocytosis of substrates. Measured by flow cytometry plotted per replicate. FS scatter. FITC/pHrodo=phagocytosis of substrate.

**Figure S4:**
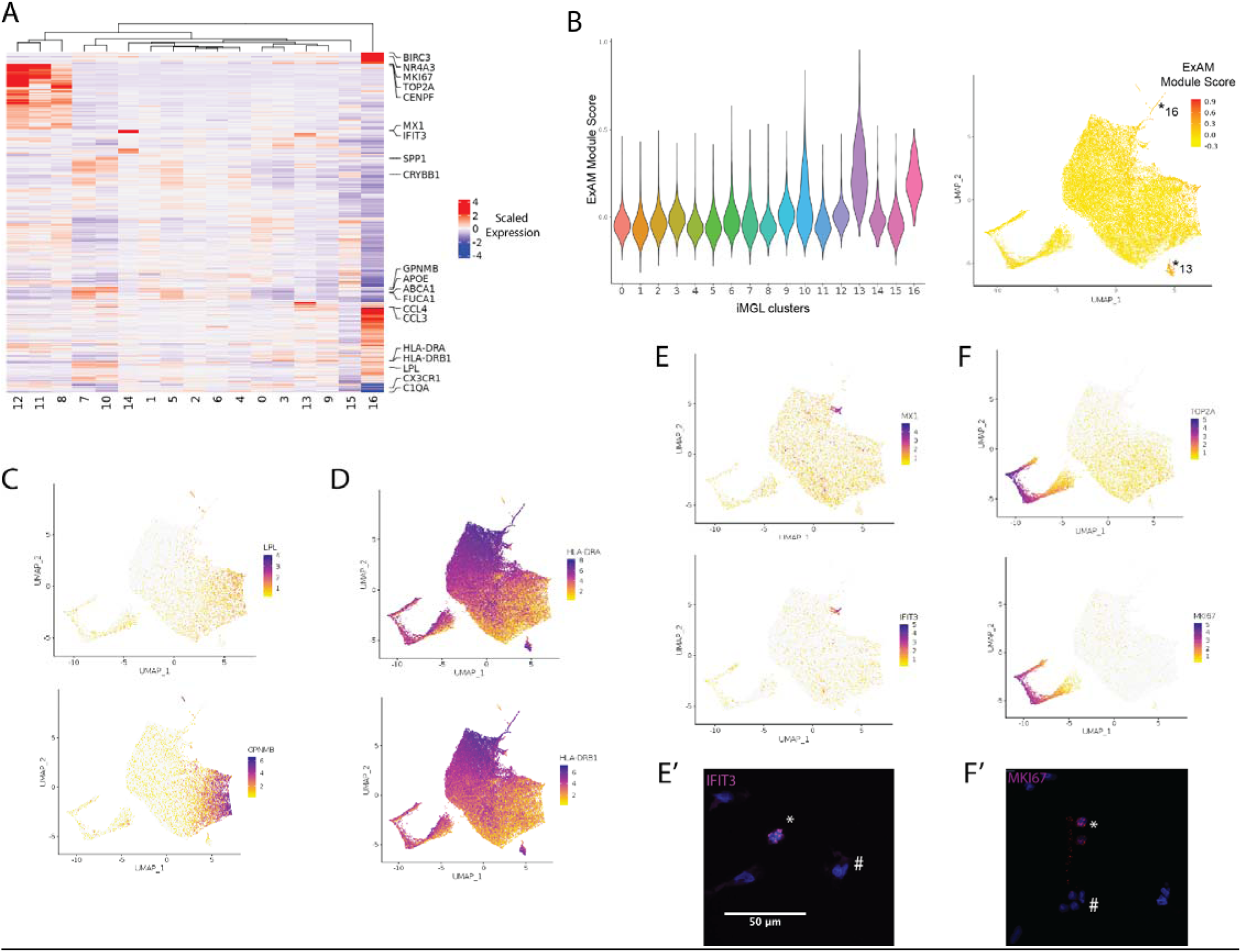
Differential expression analysis reveals distinct iMGL states. A) Heatmap of all differentially expressed genes identified per cluster. A subset of marker genes for different clusters are highlighted. B) Aberrant activation (ExAM) module score^27^ plotted per iMGL cluster (left). UMAP projection of the ExAM module score with clusters iMGL_13 and iMGL_16 highlighted. C) Expression of LPL and GPNMB (iMGL_7 and iMGL_10). D) Expression of HLA-DRA and HLA-DRB1 (marker genes for iMGL_9 and iMGL_3). E) Expression of MX1 and IFIT3 (marker genes for iMGL_14). E’) In-situ hybridization for IFIT3 reveals positive (*) and negative (#) iMGLs. F) Expression of TOP2A and MKI67 (marker genes for iMGL_8, iMGL_11, iMGL_12). F’) In-situ hybridization for MKI67 reveals positive (*) and negative (#) iMGLs.

**Figure S5:**
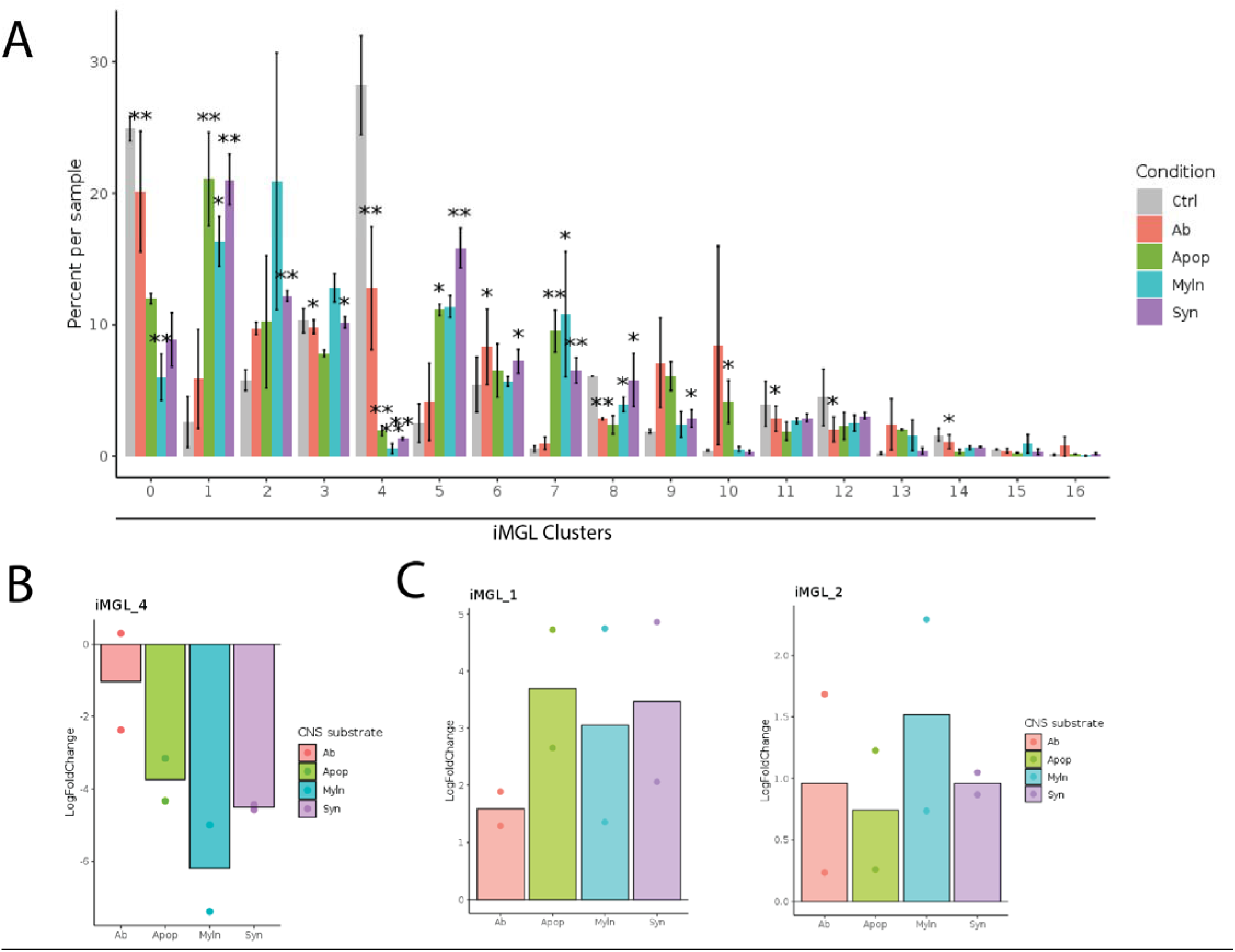
Distribution of iMGL clusters by substrate exposure condition. A) Barchart of percentage composition of cells per condition per cluster. Statistical significance determined by Dirichlet regression. * = p-value<0.05 and ** = p-value<0.01. change of iMGL_4 relative to the untreated control condition showing broad decrease in response to substrates. C) Fold iMGL_1 and iMGL_2 relative to the untreated control condition showing broad increase in response to substrates.

**Figure S6:**
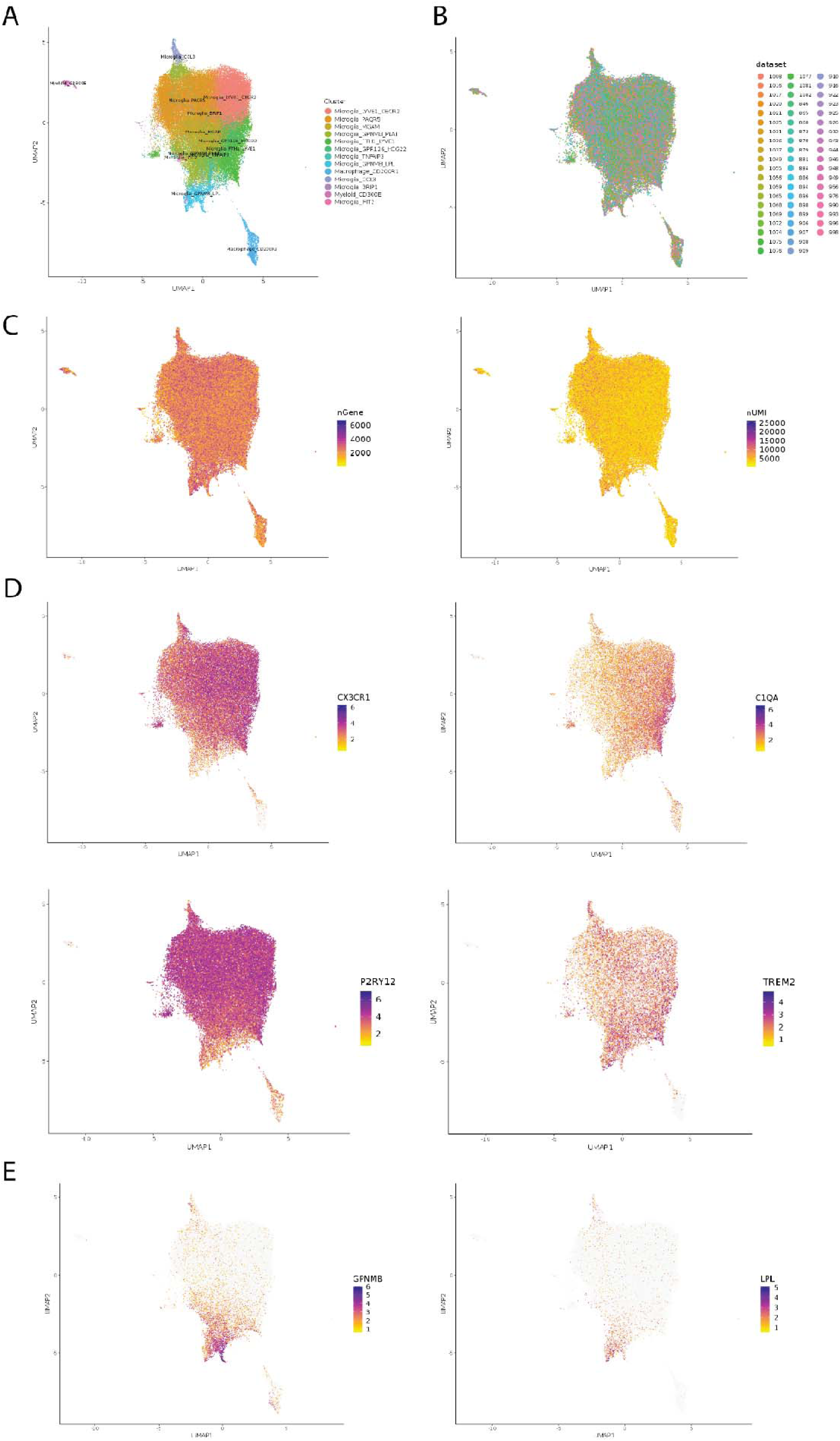
Summary of in vivo human microglial dataset used in this study: A) UMAP projection of cluster annotations of *in vivo* cortical biopsy single nucleus RNA transcriptomic dataset from Kamath et al (in preparation). Dataset consists of 58 subjects and 73,267 single microglial/myeloid nuclei. Of particular note is the Microglia_GPNMB_LPL which are human DAMs (Kamath et al. in preparation). B) UMAP projection colored by patient. No cluster consists of data from a single patient and there is good distribution of microglia across clusters. C) UMAP projection of two critical quality control metrics, number of genes (nGene, left) and number of unique molecular identifiers (nUMI, right). D) Expression of key microglial genes *CXCR1*, *C1QA*, *P2RY12* and *TREM2.* E) Expression of key DAM marker genes *GPNMB* and *LPL*^23, 24^.

**Figure S7:**
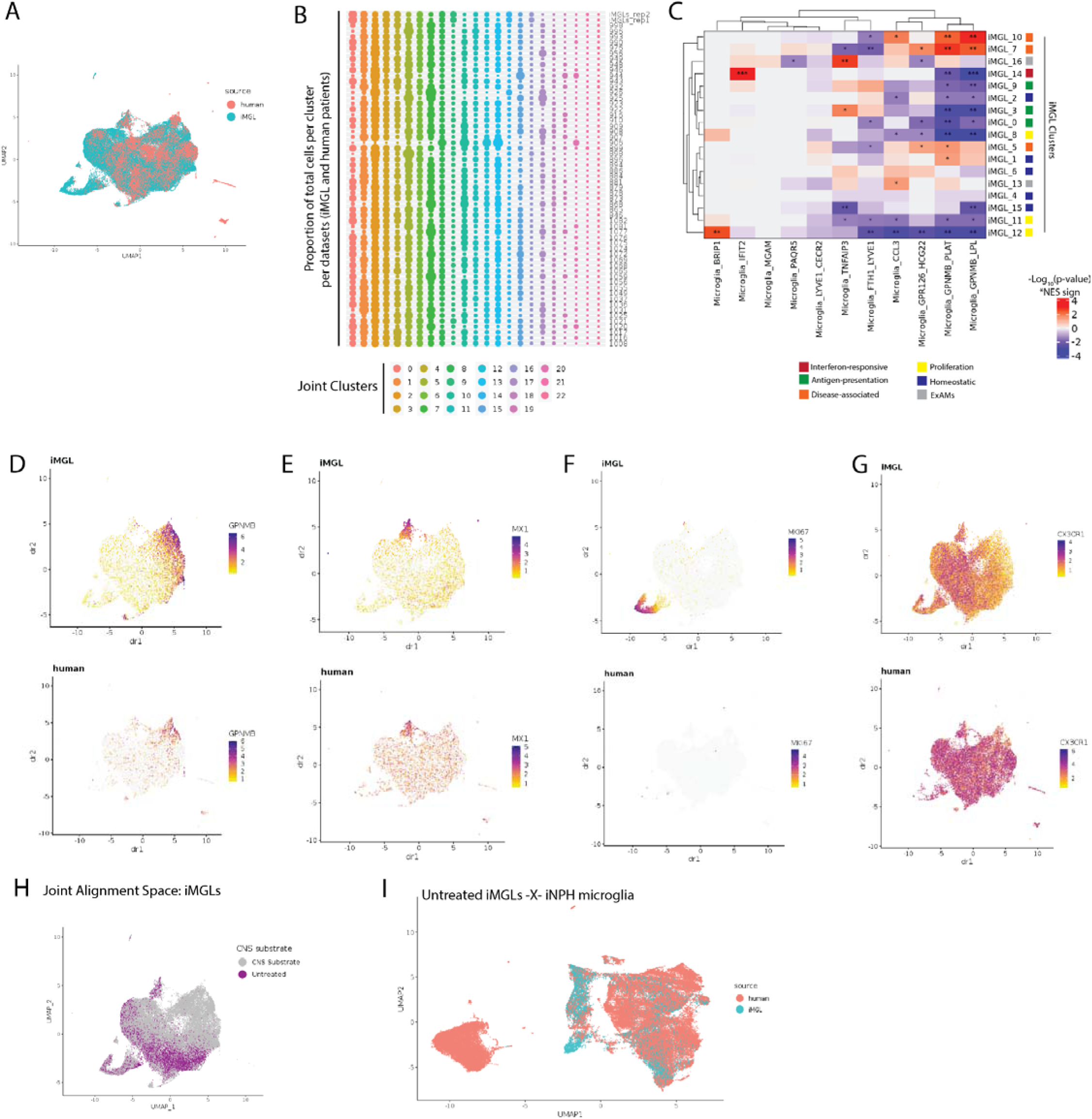
iMGL and *in vivo* human microglia dataset integration using LIGER. A) UMAP projection of iMGLs and in-vivo human cortical microglia after dataset integration using LIGER. Cells plotted by dataset source (iMGLs or human) demonstrating good integration between two data sources (alignment score = 0.76). B) Proportion of cells per joint cluster for each dataset (iMGL replicates or individual human sampled), dots are sized by proportion of total cells per dataset. C) Heatmap illustrating the relative enrichment significance (determined by geneset enrichment analysis) of positively enriched marker genes from each in vivo human cluster (from Kamath et al.) within differentially expressed genes of each iMGL cluster. D-G) UMAP projection of iMGL (top) or human (bottom) cells from the integration with expression of key marker genes *GPNMB* marker for disease-associated (D), *MX1* marker for interferon responsive (E), *MKI67* proliferative microglia (F) and *CX3CR1* homeostatic microglia (G). H) UMAP projection of iMGLs only in the joint alignment space with single cells colored by substrates or untreated demonstrating addition of substrates is key for generating transcriptional diversity. I) UMAP projection of untreated iMGLs integrated with *in vivo* human microglia using LIGER (otherwise same parameters used) results in poorer alignment (alignment score=6.25). Cells plotted by dataset source (iMGLs or human)

**Figure S8:**
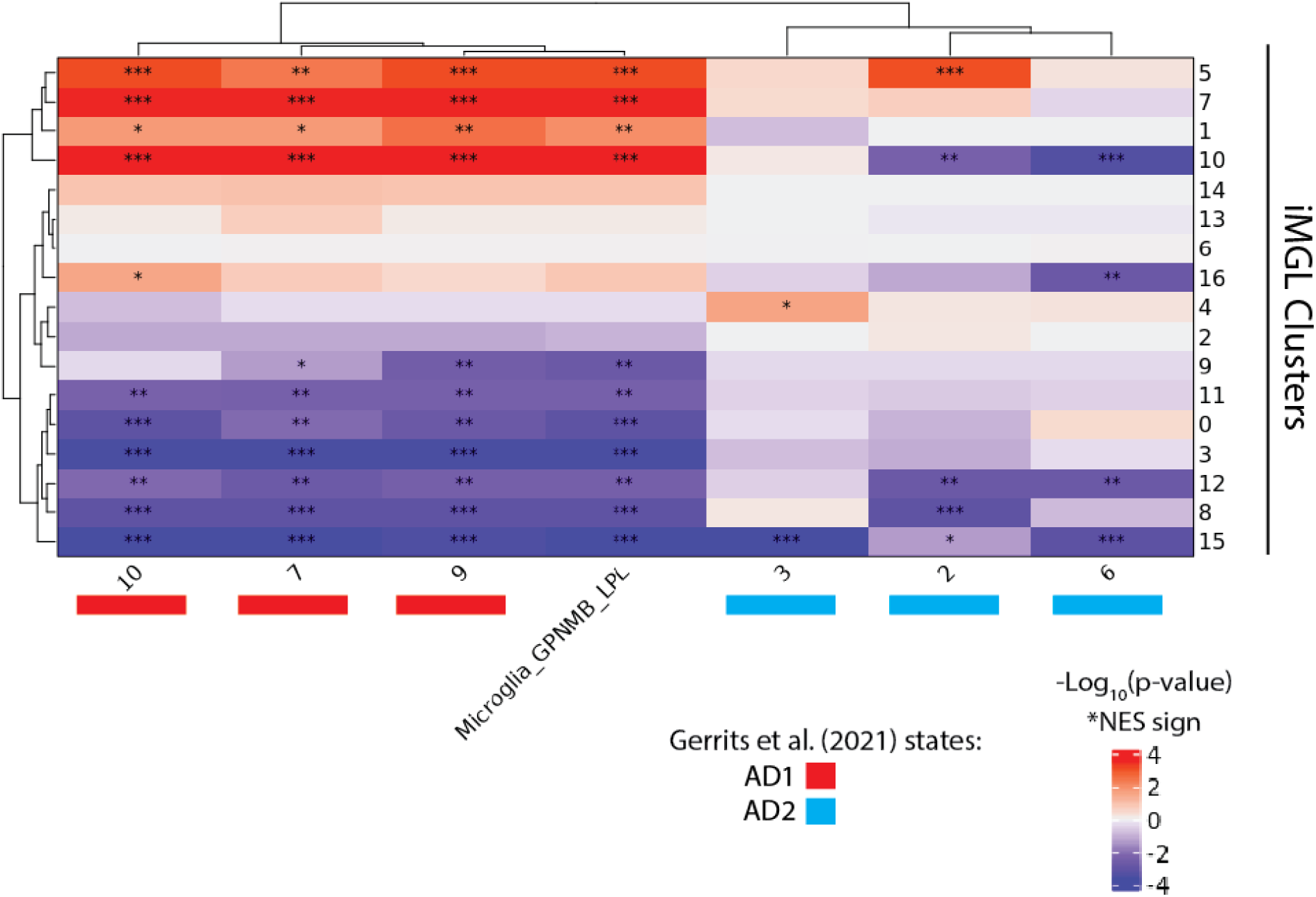
Geneset enrichment analysis confirms DAM signature in iMGLs: Heatmap illustrating the relative enrichment significance (determined by geneset enrichment analysis) of positively enriched DAM genes (derived from Gerrits et al 2021 and Kamath et al. in preparation) in differentially expressed genes for each iMGL cluster. AD1 (10, 7, 9) and AD2 (3, 2, 6) are two disease-associated states identified in-vivo as correlated with AD patient status^24^. AD1 correlates with amyloid and colocalizes with plaques while AD2 correlate with tau pathology^24^. Microglia_GPNMB_LPL are DAMs from Kamath et al, (in preparation).

**Figure S9:**
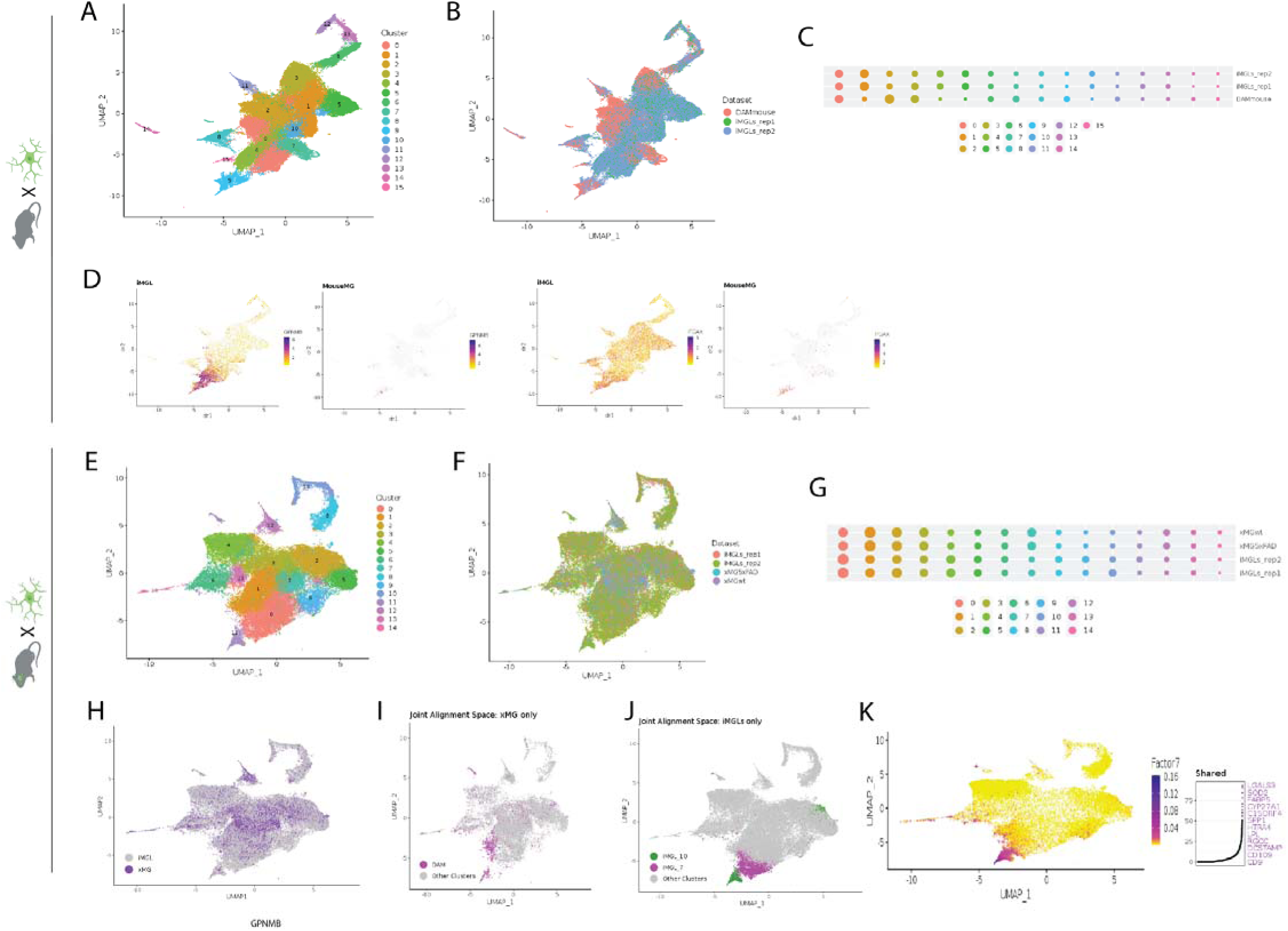
Dataset integration of iMGLs with mouse AD and xenotransplanted human microglia. A) UMAP projection of iMGLs and in-vivo mouse microglia (from wild-type and 5xFAD genotypes) after dataset integration. Cells labeled by joint clustering from this analysis. B) Cells labeled by dataset (iMGL replicate or mouse). C) Proportion of cells from either iMGLs or mouse datasets for each joint cluster in the mouse/iMGL alignment. D) UMAP plot of iMGLs (left) or mouse microglia (right) with expression of disease-associated microglia markers *GPNMB* and *ITGAX*. E) UMAP projection of iMGLs and xenotransplanted microglia (into transplantation competent wild-type or 5xFAD backgrounds) after dataset integration. Cells labeled by joint cluster. F) UMAP projection of iMGL and xenotransplated microglia (xMG) dataset integration labeled by dataset (iMGL replicates or xMGs in WT or 5xFAD mice). G) Proportion of cells from either iMGLs or xMGs for each joint cluster. H) Cells labeled by data source (iMGLs or xMGs). I-J) Plotting disease associated microglia (DAMs) in xMGs or iMGLs (iMGL_7 and iMGL_10) in the joint integration space. K) UMAP projection of LIGER alignment showing the shared factor common to iMGL and xMG DAMs. Right, top constituent genes of this shared factor.

**Figure S10:**
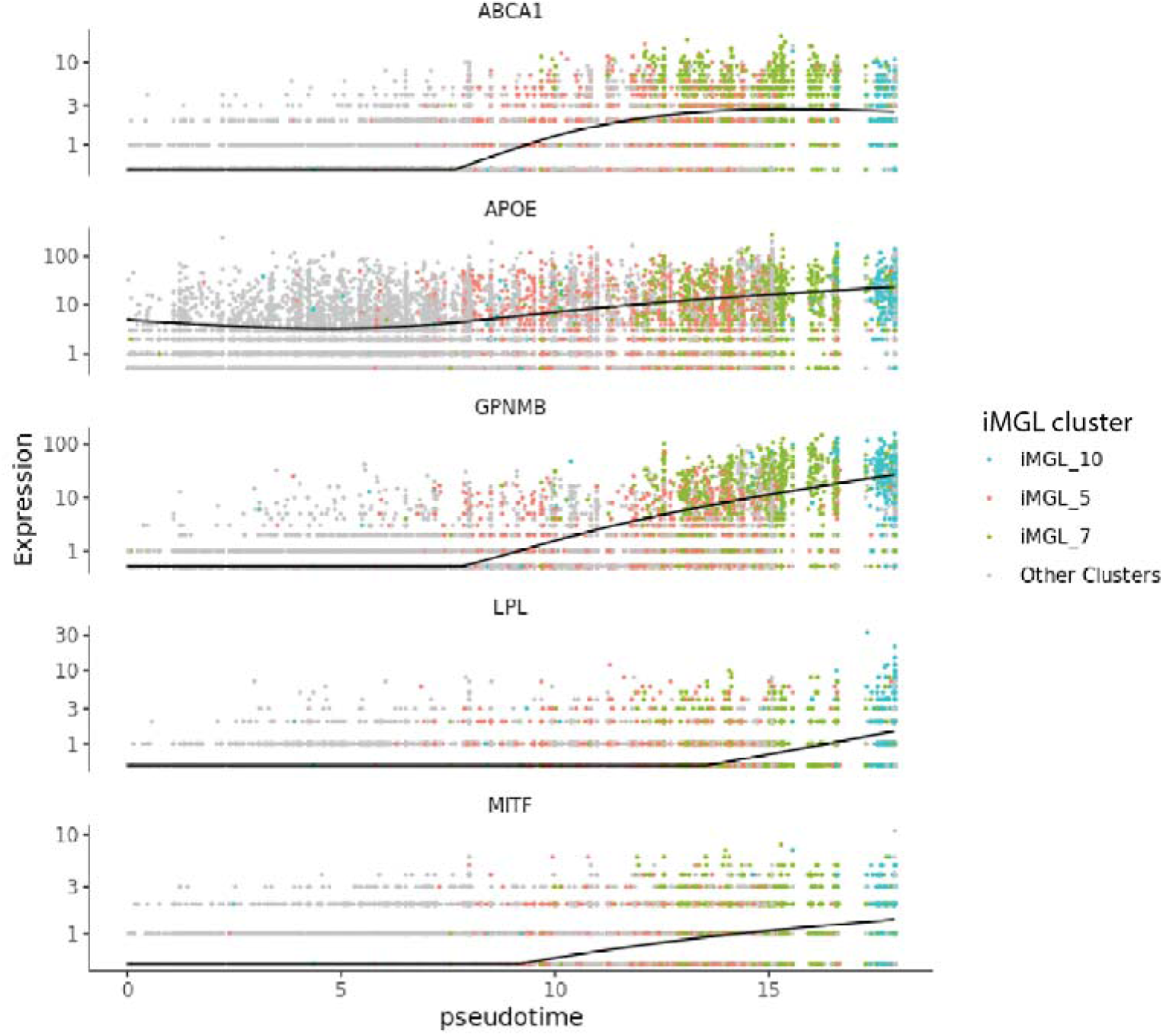
Clusters iMGL_5, iMGL_7 and iMGL_10 form a trajectory of the DAM signature: Single cell profiles of apoptotic neuron-exposed iMGLs (n=2) ordered by pseudotime and the expression of two five key human DAM genes plotted. Cells are labeled by cluster identity.

**Figure S11:**
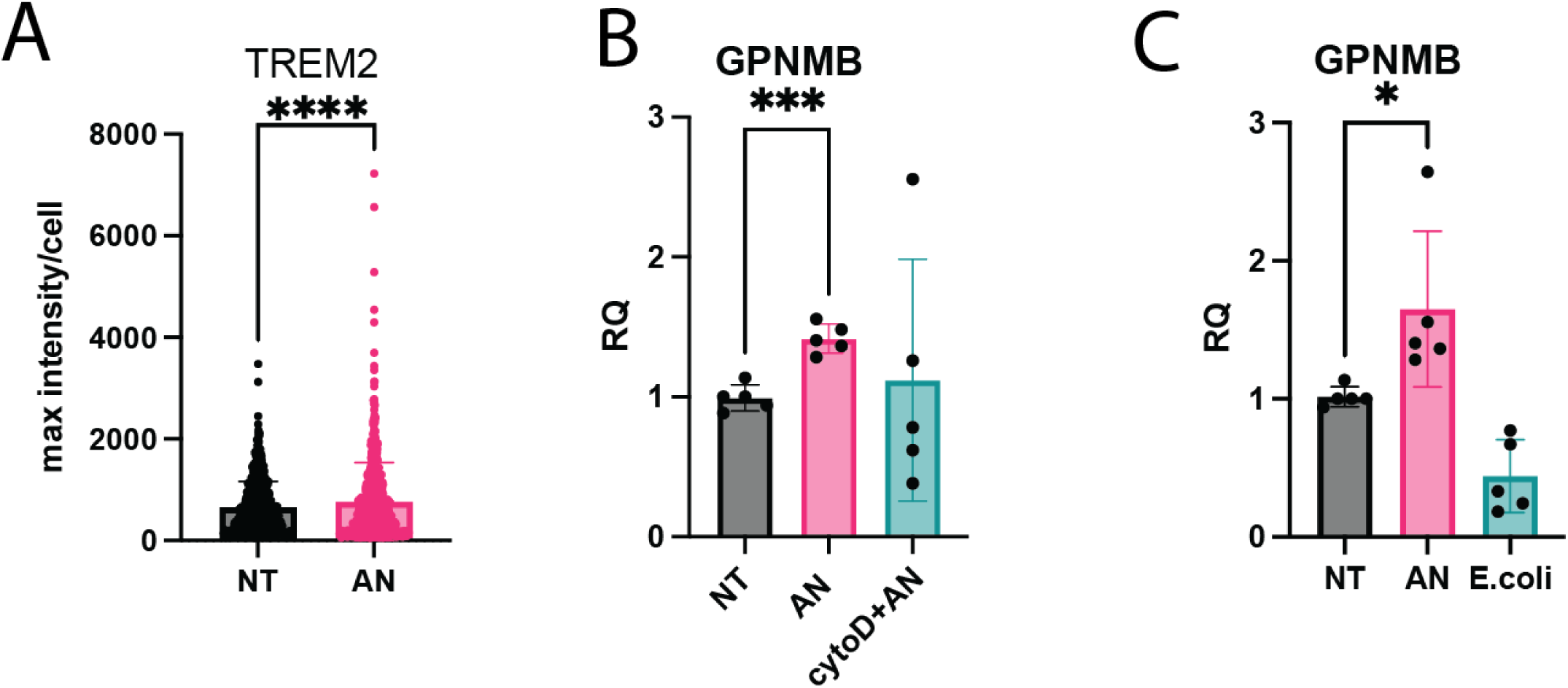
Apoptotic neurons exposure induces TREM2 and GPNMB, and is dependent on phagocytosis and neuronal substrate exposure: A) Relative intensity of TREM2 antibody stain determined by immunocytochemistry shows enrichment after exposure to apoptotic neurons (AN) compared to non-treated (NT) (p-value<0.0001). B) Induction of GPNMB mRNA in iMGLs (specific marker of iMGL_7 and iMGL_10) by exposure to apoptotic neurons (AN) is blocked by cytochalasin D (cytoD+AN), a phagocytosis inhibitor. (p-value<0.0001) C) Induction of GPNMB does not occur in response to a non-neurodegenerative substrate, E. coli. For B-C: Apoptotic neuron treatment was for 24 hours, gene expression data acquired by qPCR (p-value= 0.0374).

**Figure S12:**
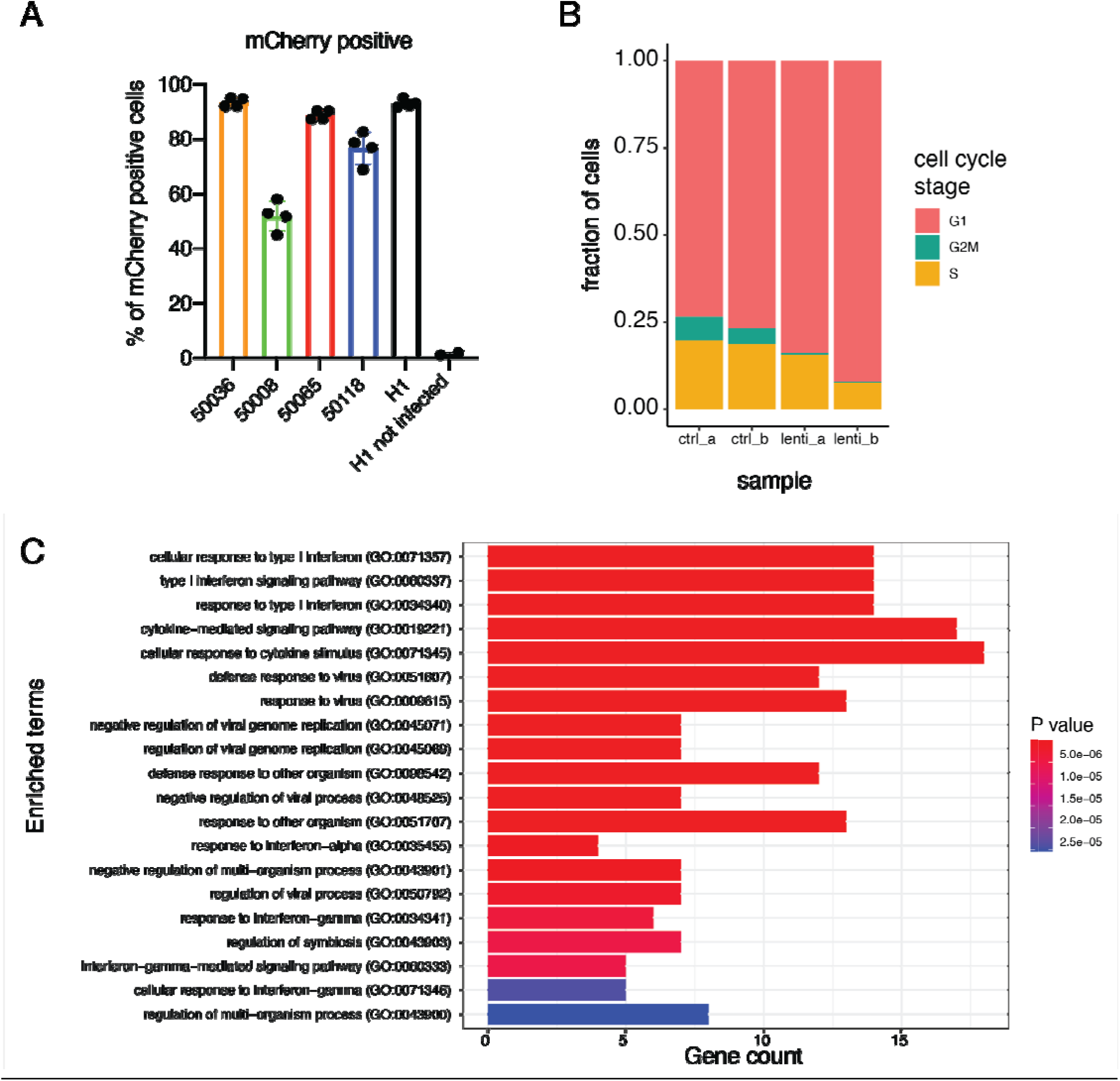
Lentiviral and VPX co-transduction enables broad transduction of iMGLs derived from different iPSC lines and characterization of interferon response. A) Transduction efficiency of iMGLs transduced from multiple cell lines using co-transduction strategy with VPX. Vector is non-targeting Cas9 control co-expressing mCherry and positive cells determined by flow cytometry. B) Distribution of cells by cell cycle stage in the control and lentivirus transduced samples. C) Gene ontology analysis of differentially expressed genes between control and lentivirus transduced samples. For this analysis, cells in the S and G2M stage of cell cycle were removed, to focus on gene expression changes other than those related to cell cycle.

**Figure S13:**
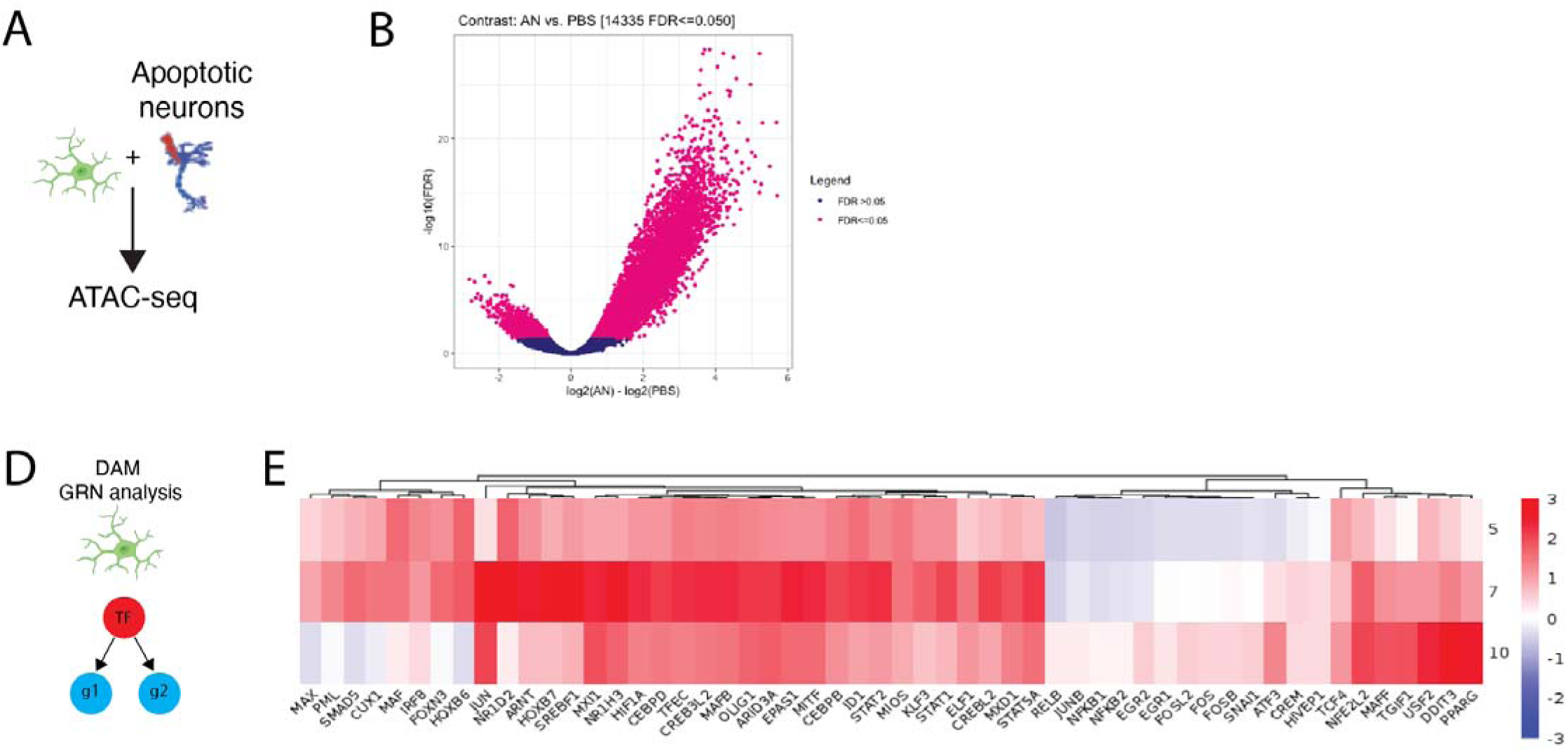
Epigenetic dissection of the iMGL response to apoptotic neuron exposure. A) Schematic of the ATAC-seq experiment. iMGLs were exposed to apoptotic neurons prior to analysis. B) Volcano plot illustrating differentially enriched peaks in the ATAC-seq data from iMGLs exposed to apoptotic neurons compared to untreated iMGLs. Statistically significant (FDR <=0.05) peaks are shown in magenta. C) Volcano plot illustrating differentially enriched peaks in the H3K27ac ChIP-seq data from iMGLs exposed to apoptotic neurons compared to untreated iMGLs. Statistically significant (FDR <=0.05) peaks are shown in magenta. D) Schematic for gene regulatory network (GRN) analysis using SCENIC. E) Identified TFs with GRNs found to be differentially enriched (see methods) in DAM iMGL clusters (iMGL_5, iMGL_7 and iMGL_10) are plotted by a scaled and centered area under the curve (AUC) for each regulon.

**Figure S14:**
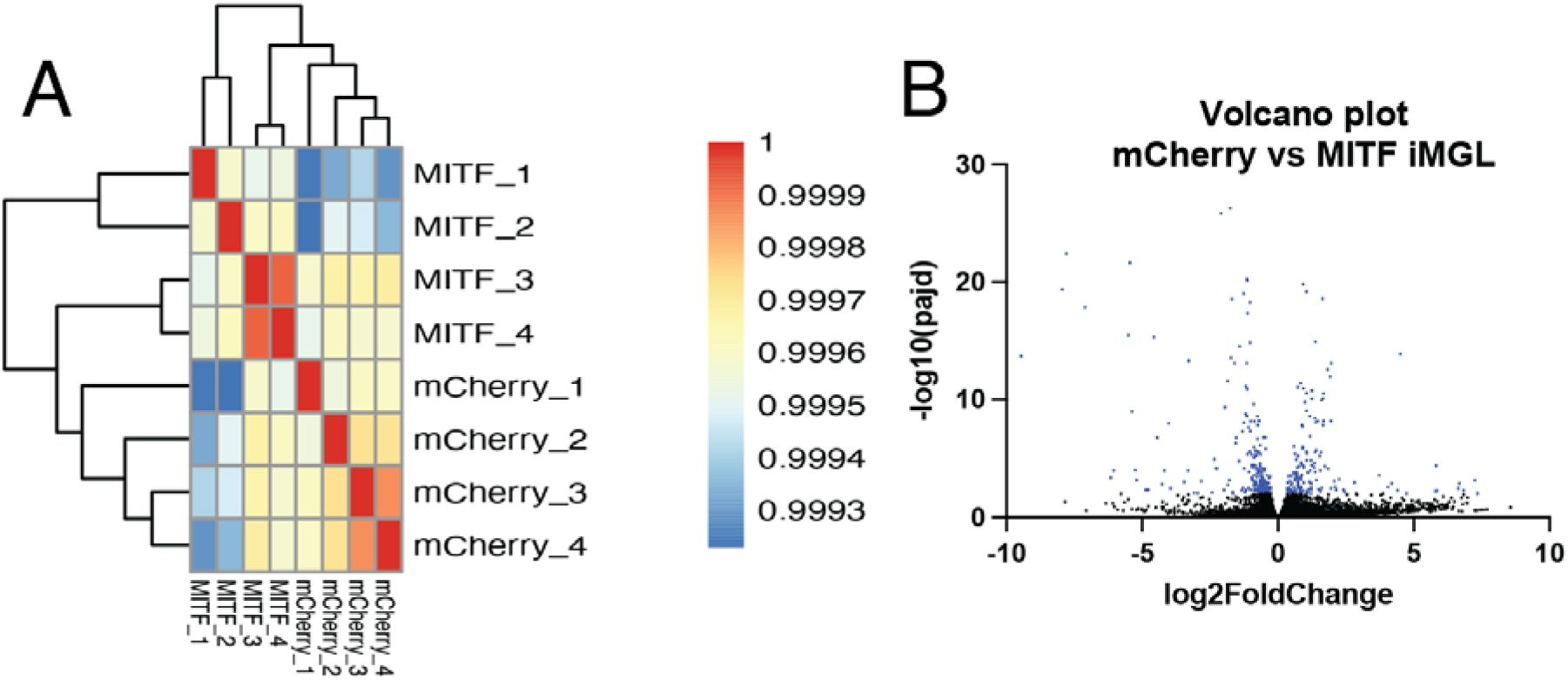
Further analysis of RNAseq data of iMGLs transduced with either mCherry or MITF. A) Correlation heatmap of RNAseq samples after MITF overexpression or mCherry expression B) Volcano plot illustrating differentially enriched genes in the iMGLs overexpressing MITF compared to iMGL overexpressing mCherry as control. Statistically significant (FDR <=0.05) peaks are shown in blue.

**Figure S15:**
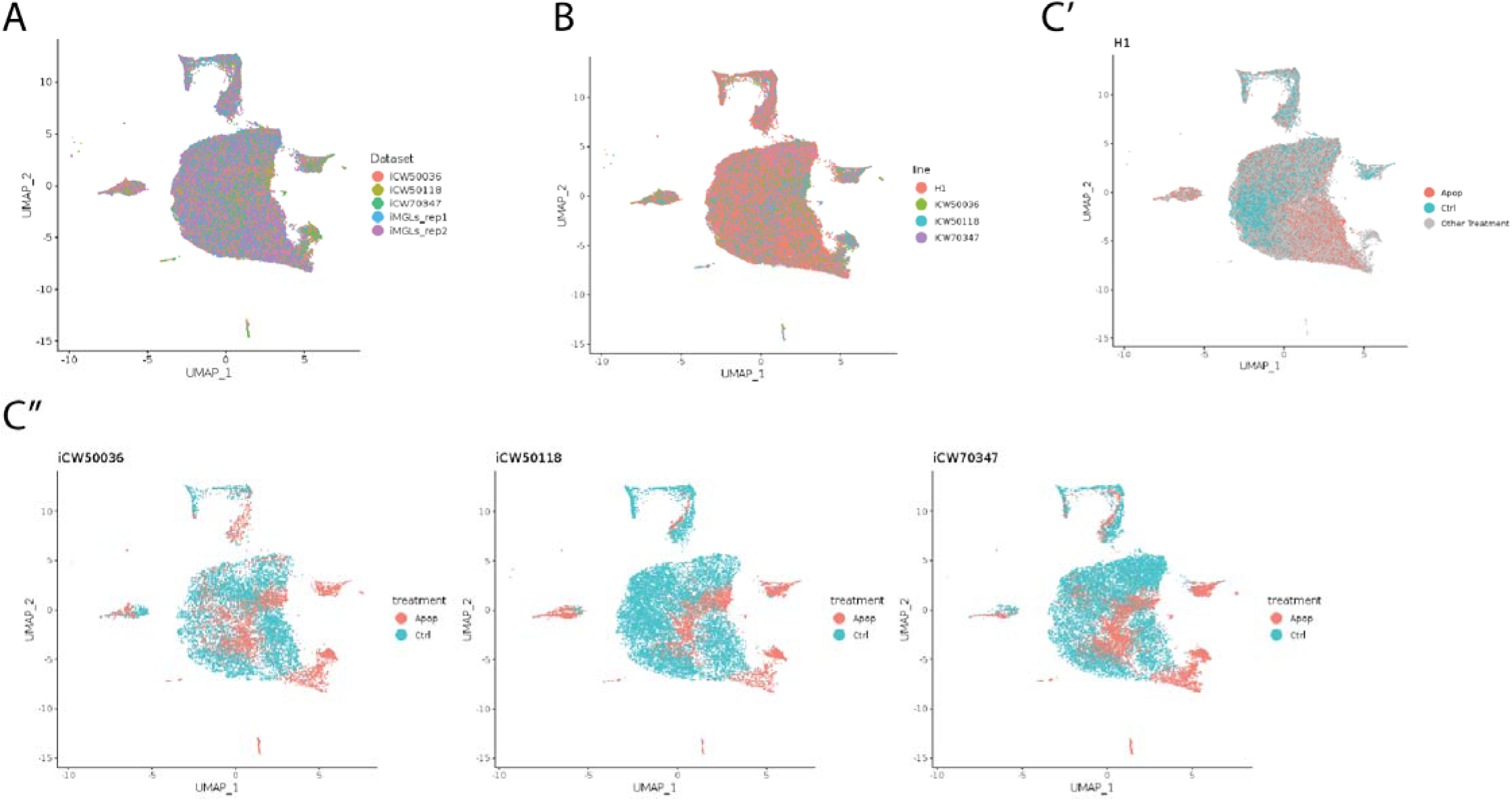
Comparison of ESC-derived and iPSC-derived iMGLs revealed similar transcriptional signatures. A) UMAP projection of LIGER dataset integration of ESC-derived (Figure 1) and iPSC-derived iMGLs reveal good integration. Cells colored by dataset. B) Same as A except cells are labeled by line. C) UMAP projection of cells plotted by dataset source and colored by either untreated (cyan) or apoptotic neuron treatment (red) shows good correspondence between H1- (C’) and iPSC-derived iMGLs (C’’).

**Figure S16:**
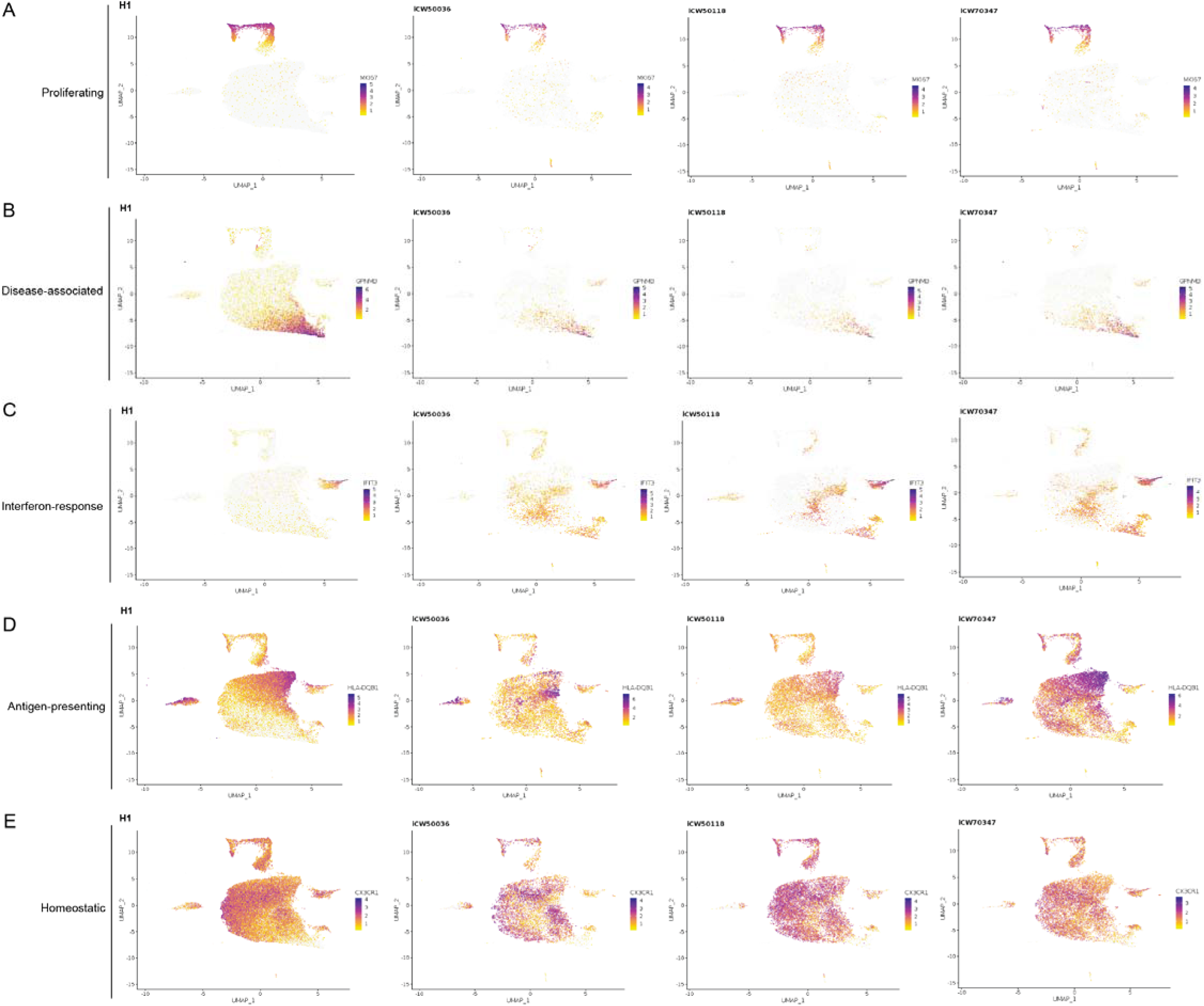
ESC-derived and iPSC-derived iMGLs share similar transcriptional states: A-E) UMAP projection of cells from the ESC- and iPSC-derived iMGL dataset integration, plotted by dataset (H1 and iPSC line), showing expression of specific state marker genes A) Proliferating, *MKI67*. B) Disease-associated, *GPNMB*. C) Interferon-response, *IFIT3*. D) Antigen-presenting, *HLA-DQB1*. E) Homeostatic, *CXCR3*.

**Figure S17:**
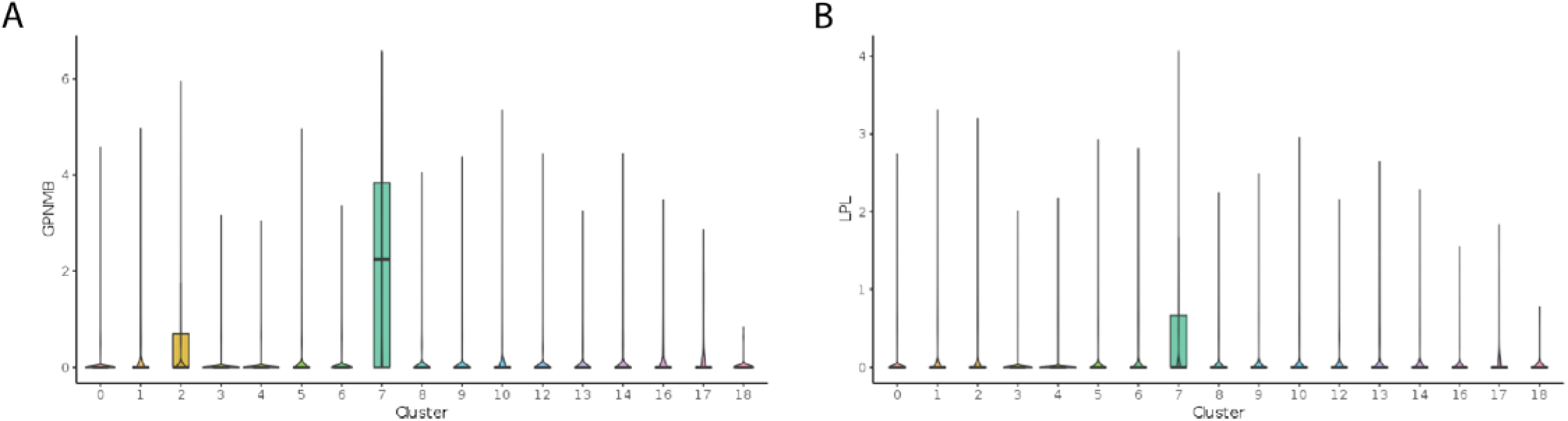
Expression of DAM marker genes in ESC-derived and iPSC-derived iMGL dataset integration: A-B) Violin plots showing expression of GPNMB (A) and LPL (B) across joint clusters from the ESC and iPSC iMGL dataset integration. Cluster 7 represents putative DAM clustering.

